# Global BioID-based SARS-CoV-2 proteins proximal interactome unveils novel ties between viral polypeptides and host factors involved in multiple COVID19-associated mechanisms

**DOI:** 10.1101/2020.08.28.272955

**Authors:** Estelle M.N. Laurent, Yorgos Sofianatos, Anastassia Komarova, Jean-Pascal Gimeno, Payman Samavarchi Tehrani, Dae-Kyum Kim, Hala Abdouni, Marie Duhamel, Patricia Cassonnet, Jennifer J. Knapp, Da Kuang, Aditya Chawla, Dayag Sheykhkarimli, Ashyad Rayhan, Roujia Li, Oxana Pogoutse, David E. Hill, Michael A. Calderwood, Pascal Falter-Braun, Patrick Aloy, Ulrich Stelzl, Marc Vidal, Anne-Claude Gingras, Georgios A. Pavlopoulos, Sylvie Van Der Werf, Isabelle Fournier, Frederick P. Roth, Michel Salzet, Caroline Demeret, Yves Jacob, Etienne Coyaud

## Abstract

The worldwide SARS-CoV-2 outbreak poses a serious challenge to human societies and economies. SARS-CoV-2 proteins orchestrate complex pathogenic mechanisms that underlie COVID-19 disease. Thus, understanding how viral polypeptides rewire host protein networks enables better-founded therapeutic research. In complement to existing proteomic studies, in this study we define the first proximal interaction network of SARS-CoV-2 proteins, at the whole proteome level in human cells. Applying a proximity-dependent biotinylation (BioID)-based approach greatly expanded the current knowledge by detecting interactions within poorly soluble compartments, transient, and/or of weak affinity in living cells. Our BioID study was complemented by a stringent filtering and uncovered 2,128 unique cellular targets (1,717 not previously associated with SARS-CoV-1 or 2 proteins) connected to the N- and C-ter BioID-tagged 28 SARS-CoV-2 proteins by a total of 5,415 (5,236 new) proximal interactions. In order to facilitate data exploitation, an innovative interactive 3D web interface was developed to allow customized analysis and exploration of the landscape of interactions (accessible at http://www.sars-cov-2-interactome.org/). Interestingly, 342 membrane proteins including interferon and interleukin pathways factors, were associated with specific viral proteins. We uncovered ORF7a and ORF7b protein proximal partners that could be related to anosmia and ageusia symptoms. Moreover, comparing proximal interactomes in basal and infection-mimicking conditions (poly(I:C) treatment) allowed us to detect novel links with major antiviral response pathway components, such as ORF9b with MAVS and ISG20; N with PKR and TARB2; NSP2 with RIG-I and STAT1; NSP16 with PARP9-DTX3L. Altogether, our study provides an unprecedented comprehensive resource for understanding how SARS-CoV-2 proteins orchestrate host proteome remodeling and innate immune response evasion, which can inform development of targeted therapeutic strategies.

## Introduction

Over a century after the Spanish flu pandemic, and despite undeniable medical advances in infectiology, modern human societies and economies are deeply shaken by the magnitude of the current worldwide SARS-CoV-2 outbreak. Unprecedented and coordinated research efforts are needed to overcome the current crisis. While we keep learning daily on COVID-19 clinical features, brute-force drug screening and therapeutic trials have not yet proven their efficacy. A fine understanding of SARS-CoV-2 pathogenesis at a molecular level is thus required to: (i) supervise and understand drug assays to combat viral mechanisms; and (ii), prepare potential future pandemics. To date, three major proteomic studies define most of our direct knowledge of SARS-CoV-2 protein interactions with host cell components^1,2,3^. These works rely on classical immunoprecipitation (IP) methods followed by mass spectrometry (MS)-based identification of physical interactors. The major limitation of IP-MS is the requirement of maintaining protein-protein interactions (PPIs) throughout the purification process. This implies using gentle lysis buffer to solubilize physically associated proteins without disrupting PPIs, hence precluding the detection of poorly soluble protein partners, or for which PPIs have low affinity. The proximity-dependent biotinylation technique (BioID)^4^ circumvents most of these caveats. Briefly, it consists of fusing an abortive mutant of *E. coli* biotin ligase (BirA R118G; or BirA*) at the N- or C-terminal end of a protein of interest. Upon addition of biotin in the culture media of living cells expressing the BirA*-tagged protein of interest, the BirA* moiety activates biotin to biotinoyl-AMP in an ATP-dependent manner. Due to the low affinity of mutated BirA* for this highly reactive species, it diffuses away and conjugates to nearby free amine groups within an estimated radius of ~10 nm^5^. Since proximal proteins are covalently biotinylated, it is no longer required to maintain stable interaction during the purification process. This enables stringent lysis conditions that can solubilize proteins from all compartments, including *e.g*. transmembrane proteins. Biotinylated species, *i.e*. proteins that have been in proximity of the bait protein in living cells, can then be readily captured on streptavidin affinity columns and identified by MS following on-bead digestion, as described^6^. Allowing the identification of PPIs through covalent labeling of proximal partners in living cells, regardless of their solubility or affinity, proximity-dependent interactomics techniques are of outstanding interest to gather biochemical insights on how the SARS-CoV-2 proteins operate to hijack host cell.

## Results

### Study overview

In our study we implemented BioID to capture the proximal interactors of the 28 SARS-CoV-2 polypeptides: (**i**) ORF1a/b, two polyproteins encoding 16 non-structural proteins NSP1-16. Notably, NSP11 was excluded from our analysis due to its small size (12 AAs); (**ii**) four structural proteins: Spike (S) glycoprotein, small envelope (E) protein, membrane (M) glycoprotein and nucleoprotein (N); (**iii**) seven accessory proteins: ORF3a, ORF3b, ORF6, ORF7a, ORF7b, ORF8, ORF9b; and (**iv**), two hypothetical proteins ORF14 and ORF10 (**Supplemental Table 1**). All viral protein ORFs were first cloned in fusion with an N-ter and a C-ter BirA* tag in pDEST FRT/TO BirA* backbones by Gateway cloning, using our pDONR SARS-CoV-2 vector collection as templates^1^. To identify human protein partners, sequences encoding each BirA-fused viral protein were then integrated into Flp-In 293 T-REx cell line genomes and, following stable cell line selection, expressed in a tetracycline-inducible manner for 24 hours in biotin-supplemented media (as described^8^). Importantly, to mimic an infectious context and trigger cellular antiviral-response resembling what occurs following SARS-CoV-2 infection, we conducted an additional set of experiments where the 28 cell lines expressing N-terminally BirA*-tagged viral genes were stimulated by poly(I:C) transfection for five hours prior biotin labeling induction. This set of experiments was designed to investigate how viral proteins could interfere with proteins expressed or activated upon viral infection since it is now well established that COVID-19 is associated with a deficient interferon (IFN) type I and III response and an exacerbated inflammatory reaction^9^. Immunofluorescence assays were conducted to assess the localizations, the proper expression and efficient biotinylation capability of the bait proteins. Biotinylated species were solubilized, captured on streptavidin columns and identified by MS (see **Supplemental Material and Methods** for details). Following background removal using a set of 30 negative control runs (a series of cell samples expressing BirA* alone covering the different experimental settings), the proximal interactomes of all 28 viral proteins, each used as both N-terminally- and C-terminally-tagged baits augmented by those obtained following poly(I:C) treatment yielded 10,267 high confidence proximal interactions by a total of 2,598 host partners (**Supplemental Table 1**).

### Network topology

The global data-driven network shape reveals a core of thirteen baits and concentrated prey proteins surrounded by fifteen bait interactomes dragged out by unique set of partners (Fig. 1). This initial observation suggests that the core viral bait proteins participate in building and organizing the cellular virus factories, whereas the peripheral viral bait proteins could hijack or impair discrete cellular functions to create a suitable environment for viral survival and progeny production. More than 90% of Core prey proteins are assigned to membranes (192 to the ER, 117 to the Golgi, 127 to vesicles and 180 to plasma membranes). These membrane-rich organelles appear as the primary location of 15 viral proteins. Therefore, to facilitate data interpretation, we considered the most connected interactors as compartment-specific proximal background. To segregate specific sets of interactors between the core bait proteins, we chose to filter out all prey proteins connected to eight or more bait proteins.

**Figure 1.**
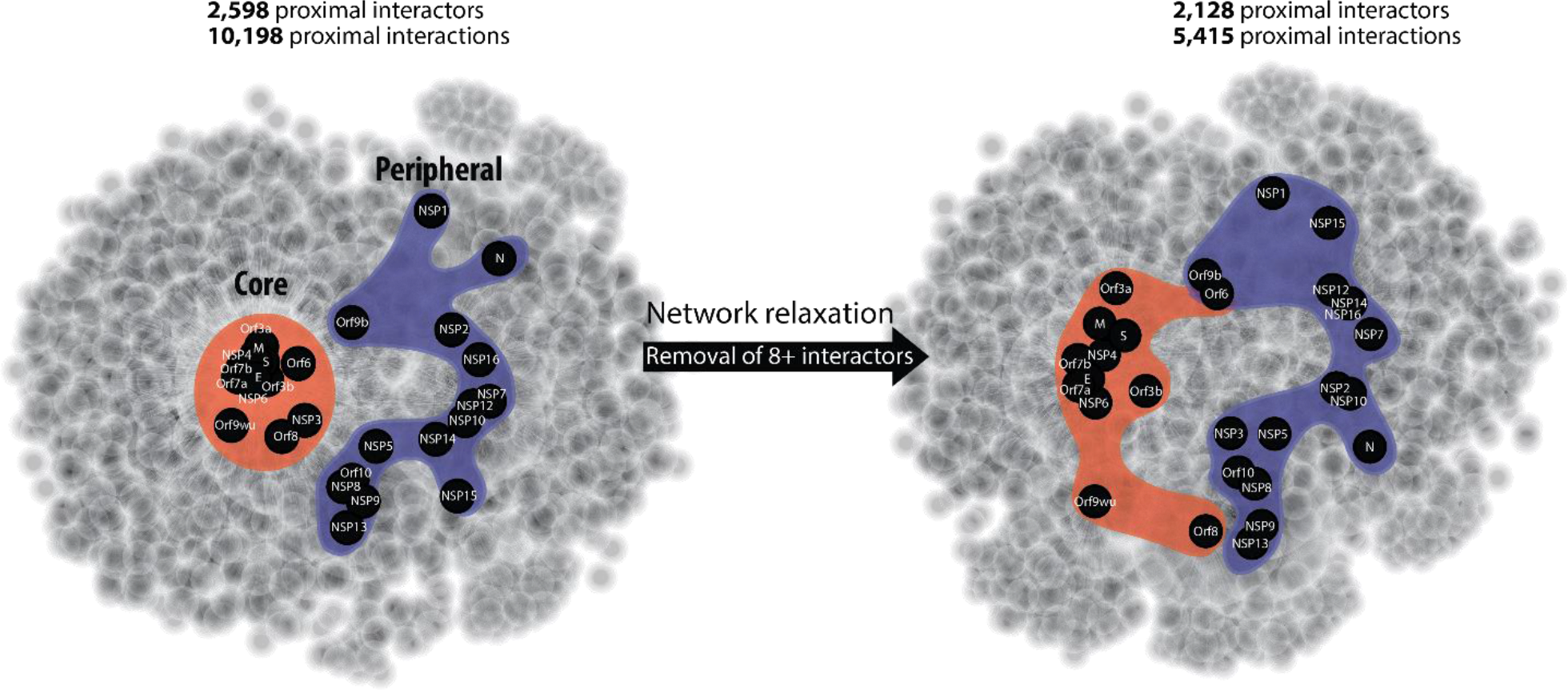
Data-driven BioID network of SARS-CoV-2 proteins. Baits and preys are positioned according to their interaction profiles; left: all high confidence proximal interactions (FDR<0.01); right: high confidence proximal interactors detected by 7 or less bait proteins.

This arbitrary filter reduced the number of interactors from 2,598 to 2,128, and the total number of interactions dropped from 10,198 to 5,415 (**Supplemental Table 1** and online dashboard). Expectedly, the core was the most affected region of the network structure. This filtering allowed its relaxation (**Fig. 1**), revealing the connection of highly enriched groups of interactor/protein complexes to specific viral bait or subsets of viral bait proteins. For each viral protein, we summarized the main locations of their interactors (**Fig. 2.A and B**). In addition to gene ontology and pathway enrichment analysis (**Supplemental Table 1; Supplemental Table 2 and Supplemental Table 3**), we manually curated the data for each viral bait proteins and assigned them to main predicted functions based on their most notable interactomes features (**Fig. 2.C**). Since our experimental design consisted in expressing a single BirA*-tagged viral protein at a time, we describe each proximal interactome profile separately. For each bait protein (**Figs. 3-31**), a Venn diagram shows the hit distribution between the different experimental settings. Given the complexity of the observed network, we reasoned that the ability to interactively explore the data would be superior to static networks, and implemented an online interactive 3D map (http://www.sars-cov-2-interactome.org/) which can be used to identify network regions, the distribution of enriched categories as well as isolate subgroups of interactions or interactors. Considering that BioID bait proteins label proximal partners within a sphere of ~10 nm radius in living cells, we argue that a data-driven network topology in three-dimensional space is an excellent representation for capturing viral protein and host factors organization. We further hypothesize that our analysis defines cellular volumes populated by identified proteins which are at the core of viral-host interplay.

**Figure 2.**
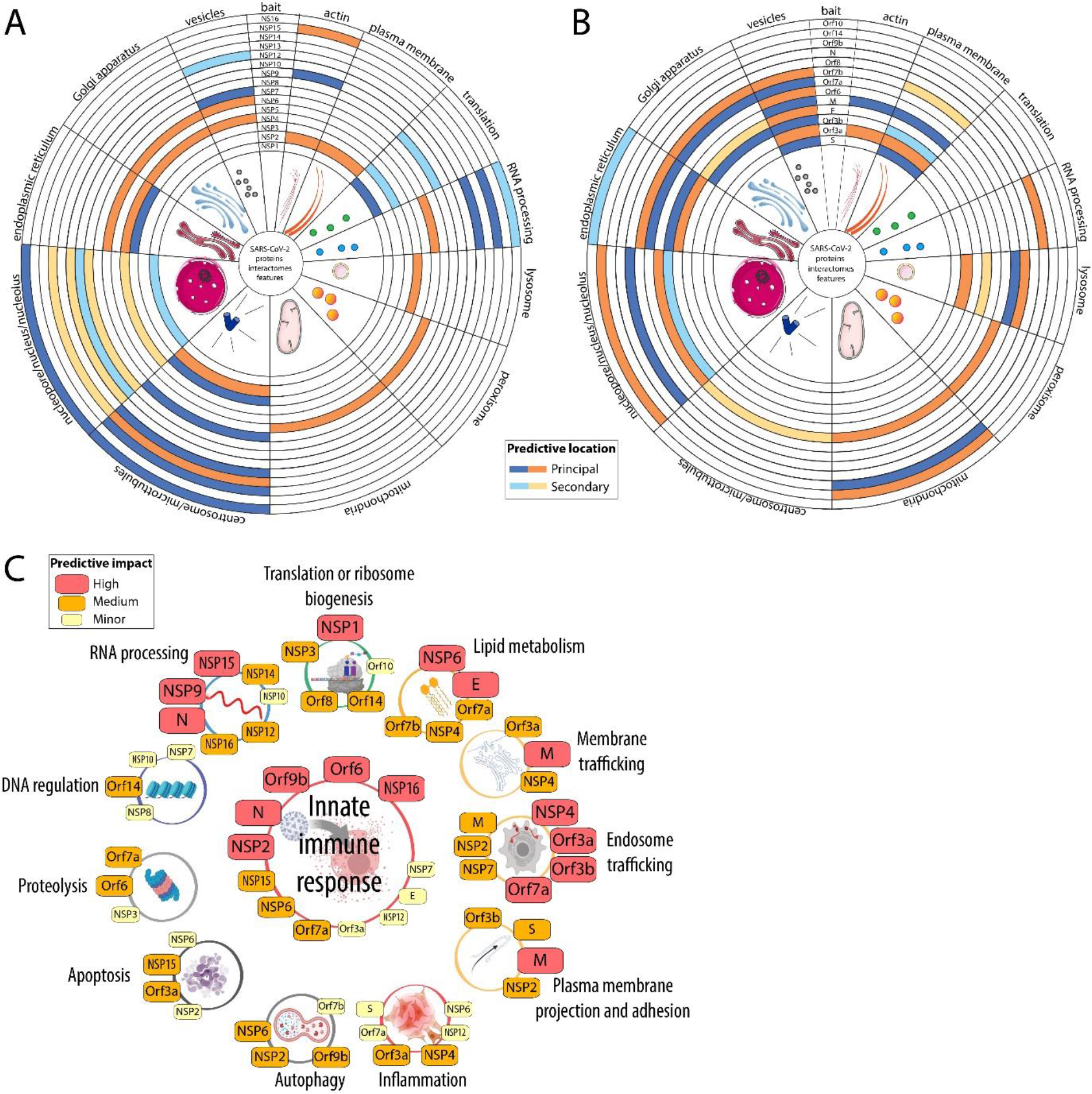
Study results overview. **A.** Probable location of the SARS-CoV-2 Non-structural proteins based on a combination of database enrichment analysis and manual curation of each bait high confidence proximal interactome. **B.** Same as (A) for the structural and accessory proteins. **C.** Assignment of probable involvement of SARS-CoV-2 viral proteins based on manual curation of their proximal interactomes. The degree of likeliness of a given viral protein to be involved in the corresponding mechanisms is color and size-coded (see legend). For each bait viral protein, the detailed analysis is following in the manuscript.

**Figure 3.**
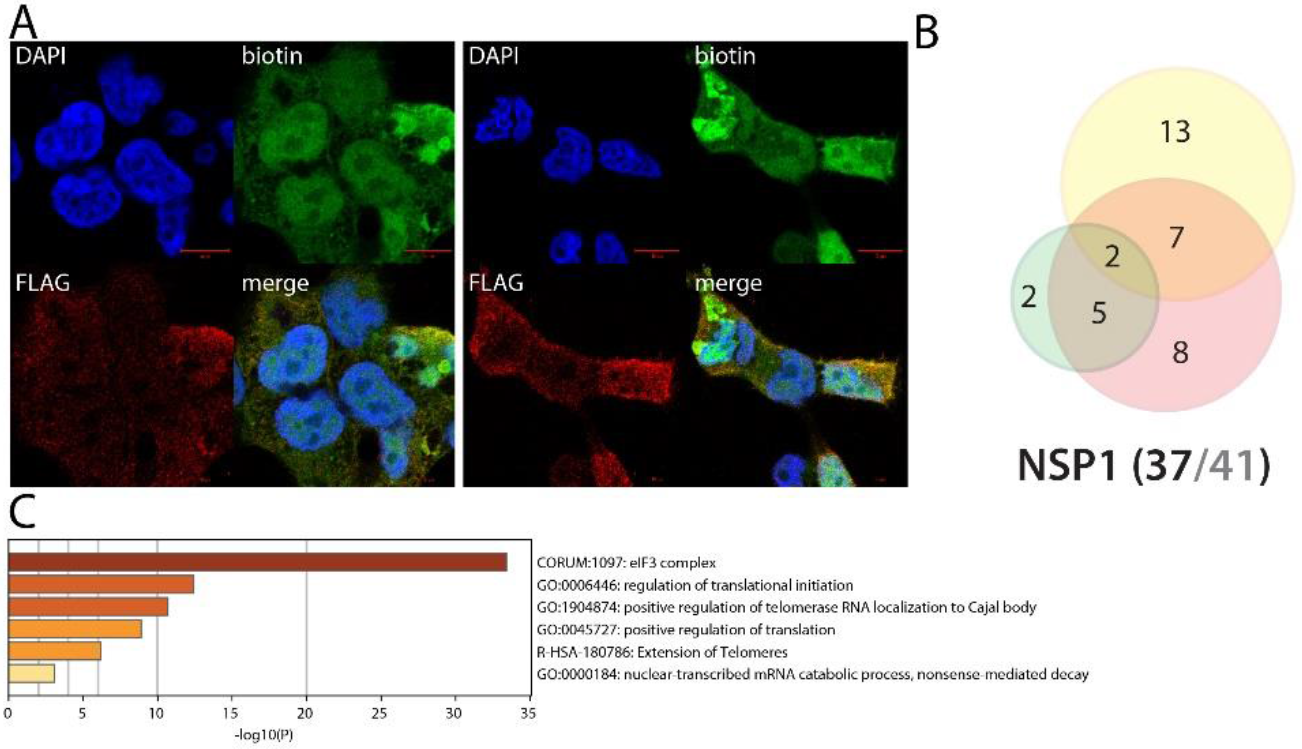
NSP1 summary. **A. (left panel)**: Confocal imaging of NSP1-BirA *Flag-expressing 293 FlpIn TREx in basal culture condition (1μg/mL of tetracycline + 50μM biotin for 24 hours); DAPI, biotin (streptavidin-AlexaFluor488; proximal interactors) and anti-FLAG (anti-mouse AlexaFluor647; bait protein) staining. **(right panel)** same staining with a pretreatment of 2μg/mL poly(I:C) before induction. **A** and **B**. Red scale bars = 10μm. **B**. Venn diagram of high confidence proximal interactors identified with the N-ter (yellow circle) and C-ter (green circle) tagged NSP1; the red circle depicts the proximal interactors detected after poly(I:C) treatment; isoforms have been collapsed to single gene names and the bait has been removed for comparison; **(bottom)** bait protein name and corresponding number of high confidence proximal interactors connected by seven or less bait proteins (bold black font) over the total number (no filtering; grey font) **C**. Top enrichment categories detected analyzing all high confidence interactors (metascape.org). The shades of orange depict -log10(P) thresholds; darker is more significant.

Together, our data provide an unprecedented coverage of the SARS-CoV-2 proteins interplay with the human cellular machineries, revealing a complex nexus of interactions that could serve as targets to counteract the infectious cycle.

### NSP1 uniquely interacts with the eukaryotic translation initiation 3 complex

Apart from a few amino-acid substitutions, SARS-CoV-1 and SARS-CoV-2 NSP1 are highly similar, and they are expected to ensure comparable functions. SARS-CoV-1 NSP1 has been reported to promote host mRNA degradation and to inhibit translation^10^. Upon SARS-CoV-1 infection, NSP1 thus appear as an important virulence factor, since it is very likely to provide the type I IFN response mRNA translation blockade^10^. In addition to previously reported interactions (*e.g*. PKP2, POLA1 and POLA2^1^) that are not involved in translation, we identify 12 components of the eukaryotic translation initiation (eIF) 3 complex uniquely associated with SARS-CoV-2 NSP1. SARS-CoV-1 NSP1 has been reported to block translation through interacting with the 40S ribosomal subunit^11^, of which we identified RPS10 as a high confidence interactor. In the previous interactomic studies, very few SARS-CoV-2 NSP1 interactors were identified^1,2,3^. Our data strongly suggest that NSP1 binds to the 40S subunit through the eIF3 complex, which is well described to modulate translational activity^12^. Besides interacting with the 40S associated proteins, we detected the RNA helicase DDX3X which has been involved in innate immune response activation in HBV^13^, VACV^14^, HCV^15^ and HIV-1^16^ infections. Of note, we observed striking toxicity expressing N-ter BirA* tagged NSP1, but not with the C-ter tagged version. Data obtained with the N-ter tagged NSP1 samples are thus only produced through transient transfection, since we were unable to generate a stable cell line for this construct. In line with this observation, the C-ter tagged version did capture only 2/12 eIF3 complex components, suggesting that NSP1 is binding to this complex mainly through its C-ter end, and that an excessive binding of NSP1 to eIF3 complex is detrimental for the cell. Under poly(I:C) treatment, NSP1 interactome was slightly modified. It gained a few interactors with no obvious involvement in viral life cycle.

### NSP2 uniquely interacts with RIG-I and STAT1 and appears as a central protein in SARS-CoV-2 innate immune response evasion

Little is known on coronaviruses NSP2 proteins functions. In our analysis, we retrieved 43 (20 in SARS-CoV1/2 and 23 with other coronaviruses) reported interactors, out of 146 total proximal proteins, including STAT1 (MHV NSP2 in mouse by BioID^17^). Only four SARS-CoV-2 interactors were overlapping with our dataset (STOML2, FKBP15, RAP1GDS1 and PCM1^1,2^), but their link to viral life cycle is yet to be determined. Our analysis revealed that NSP2 interacts with *e.g*. 28 components assigned to the cytoskeleton organization center, 34 components of plasma membrane bounded projection, a primary site of virion production^18^, and 28 components of vesicles. In addition, NSP2 BioID captured several components of the innate immune response pathways (AZI2, ZFAND5, NUMBL, USP15, STAT5B, IBTK, CYLD, TRIM26), autophagy key proteins (WIPI1, WIPI2, MAP1LC3B) and major regulators of apoptosis (*e.g*. AVEN, TP53BP2, AREL1, BAG3, ZAK, CASP8). Multiple kinases were also identified (see^18^), suggesting that NSP2 is an important signaling perturbator in infected cells. NSP2 could thus play a role in modulating cellular response to infection and death signals.

**Figure 4.**
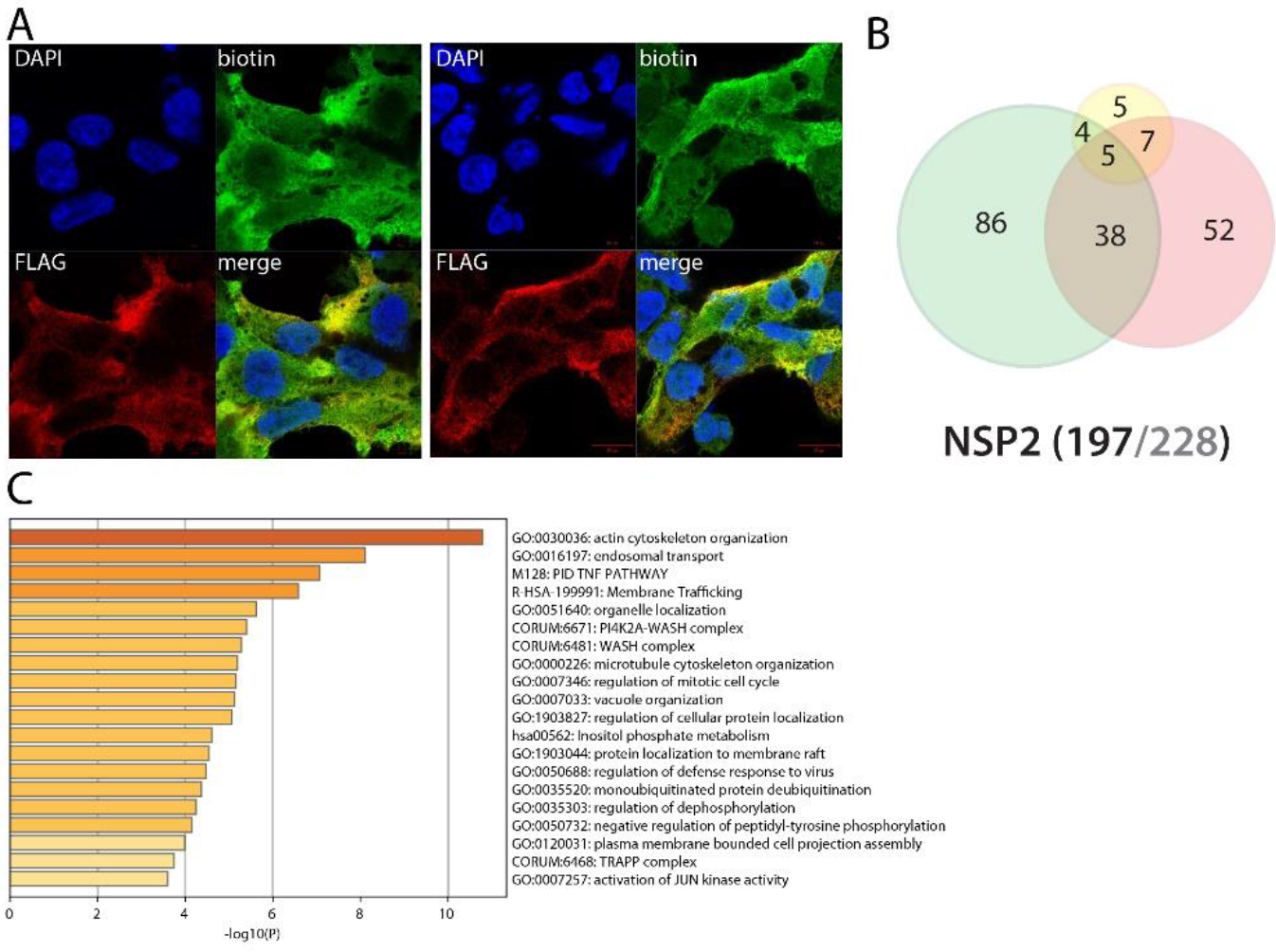
NSP2 summary. See legend Figure 1 (imaging of BirA *Flag-NSP2 here).

Upon poly(I:C) treatment, the NSP2 interactome was dramatically rewired. About 90 interactors were enriched/gained under infection-mimicking conditions, including several PPIs connecting NSP2 to major components of the antiviral response pathways. We thus uncovered the viral RNA sensor RIG-I (DDX58)^19^, the IFN response enhancing PARP9, which belongs to the ubiquitin ligase complex with DTX3L^20^, the key IFN pathway regulators IFIT1/2/3/5^21^, the signal transducer STAT1^22^, the ubiquitin ligase TRIM56^23^ and the E3 ISG15 ligase HERC5^24^ as novel NSP2 interactors. NSP2 thus appears as a core modulator of host cell innate immune response.

### NSP3 (PLpro) interacts with translation control factors and ERAD pathway components

The papain-like protease is a cysteine protease which recognizes and cleaves the LXGG motif found between NSP1 and NSP2, NSP2 and NSP3, and NSP3 and NSP4^25^. *In vitro* assays of SARS-CoV-1 PLpro also showed a deubiquitinase activity and an ability to remove ISG15 modification from substrates^26,27^. It thus appears both crucial for Orf1a viral polyprotein processing and immune evasion. PLpro is a membrane associated protein, which explains why we did detect only two overlapping interactors with classical IP-MS previous reports (MKRN2, RCHY1). Several novel interactors linked to translation control have been uncovered, *e.g*. PPP1R15A/B, LTN1 and ZNF598, EIF4E2 and GIGYF2. Besides its function towards viral polyprotein, our data suggest that NSP3 also associates with microtubule cytoskeleton dynamics (*e.g*. SLAIN1, TRIM3, KIF4, MAP7D3). Interestingly, poly(I:C) treatment induces its interaction with several components of the endoplasmic reticulum-associated degradation (ERAD) pathway, such as DERL2, HERPUD1, VIMP, and ERLIN1. Through these interactions, PLpro might regulate proper control of polyprotein translation, folding, transport and stability.

**Figure 5.**
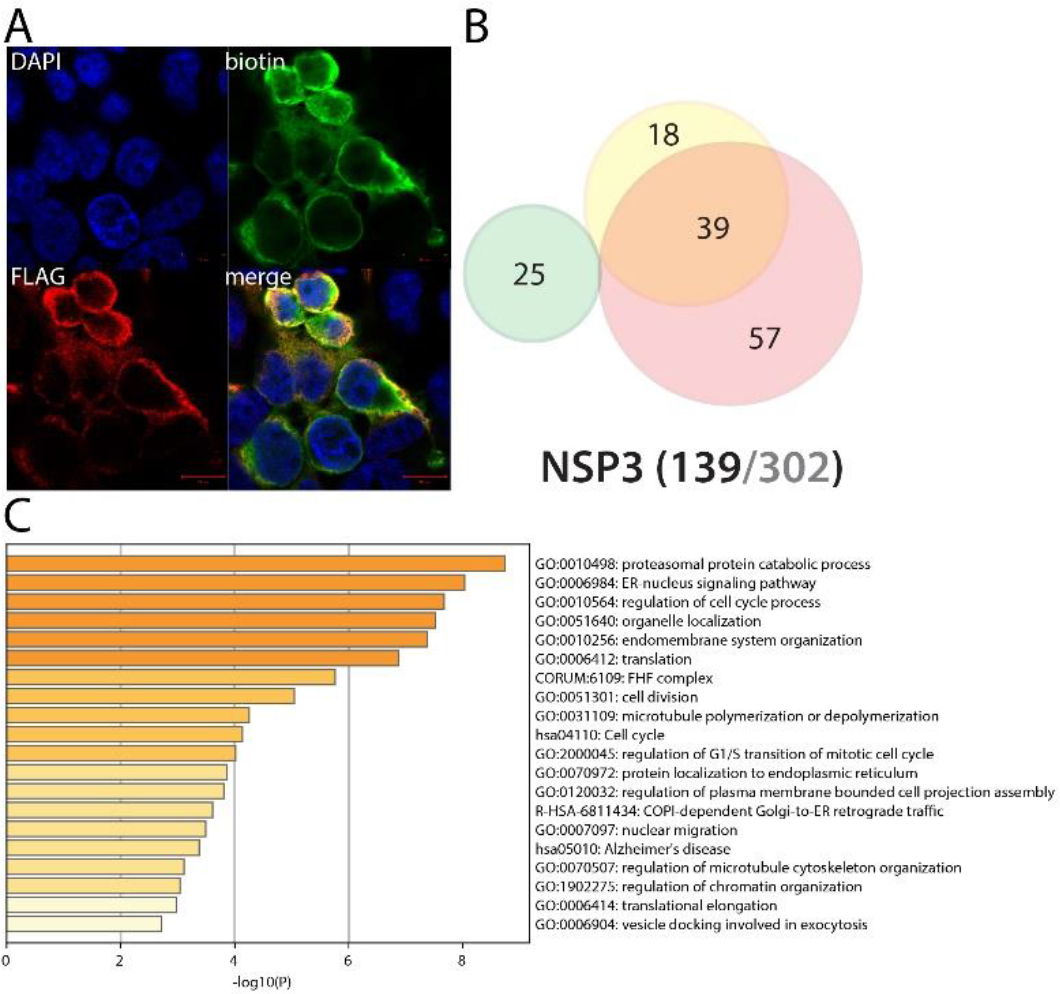
NSP3 summary. See legend Figure 1 (imaging of BirA *Flag-NSP2 here; no poly(I:C) condition).

### NSP4 interacts with endomembrane proteins and could dysregulate the endocytic degradation pathway through ITCH, MIB1 and ESCRT-0 complex upon infection

NSP4 is a transmembrane glycoprotein able to form double-membrane large vesicles associated with the SARS-CoV-1 replication complex^28^. As shown in **Fig. 6**, N-ter BirA*-tagged NSP4 alone can recapitulate resembling phenotype, suggesting that our tagging does not impair its ability to form such structure. In line with previous report^29^, we identified nuclear envelope interactors, mitochondrial associated factors and ER factors. Of note, given the membranous nature of NSP4, the BioID approach identified many novel membrane-associated interactors compared to classical IP-MS techniques. Our rich NSP4 interactome data revealed associations between *e.g*. NSP4 and the Golgi apparatus membrane, numerous vesicular complexes (*e.g*. TRAPP, SNAREs, ESCRT-0), organelle membrane contact site components and factors involved in lipid metabolism. These data suggest that NSP4 could also be involved in regulating double-membrane structure positioning and contact with multiple endomembranes. Upon addition of poly(I:C), NSP4 gained interactions with additional vesicular transport components (intra Golgi and retrograde Golgi-to-ER traffic, COPI-mediated anterograde transport), and several hits of particular interest, *e.g*. PRKCQ (Protein kinase C theta type; involved in NFκB signaling^30^), ITCH (E3 ubiquitin-protein ligase Itchy homolog) and MIB1 (Mindbomb E3 ubiquitin protein ligase 1). ITCH is a major regulator of inflammatory and antiviral pathways. It has been *e.g*. reported to trigger MAVS degradation^31^ and control NOTCH1 degradation^32^. Interestingly, it also ubiquitinylates the chemokine receptor CXCR4^33^, and directs it towards the degradative endocytic pathway targeting HGS and STAM (ESCRT-0 complex components). MIB1 is reported to interact with Notch receptor and regulates the internalization of Notch ligand. NSP4, through interacting with trafficking machineries, appears as a main host membrane remodeler and could also impair membrane embedded protein homeostasis upon SARS-CoV-2 infection.

**Figure 6.**
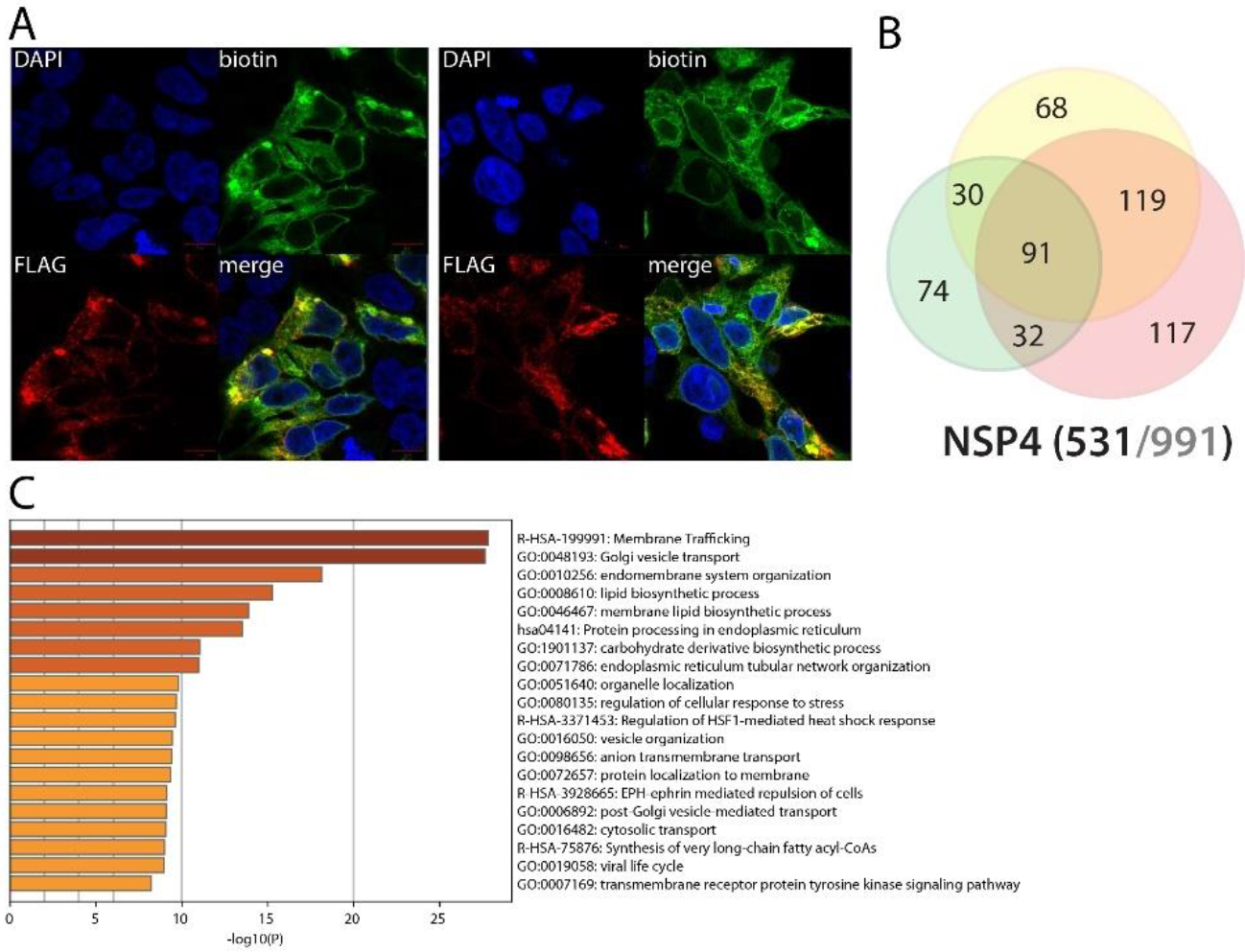
NSP4 summary. See legend Figure 1 (imaging of BirA *Flag-NSP4).

### NSP5 (3CLPro) BioID did not identify interactors enriched in specific mechanisms

The NSP5 protease (3CLPro) is the main factor processing the Orf1a/b-encoded polyproteins^34^. It is therefore an essential protein for virus protein maturation. However, our analysis only identified 14 high confidence NSP5 interactors, and we did not observe noticeable or shared features amongst these interactors. For an unknown reason, BirA*-tagged NSP5 did not efficiently capture its proximal partners.

**Figure 7.**
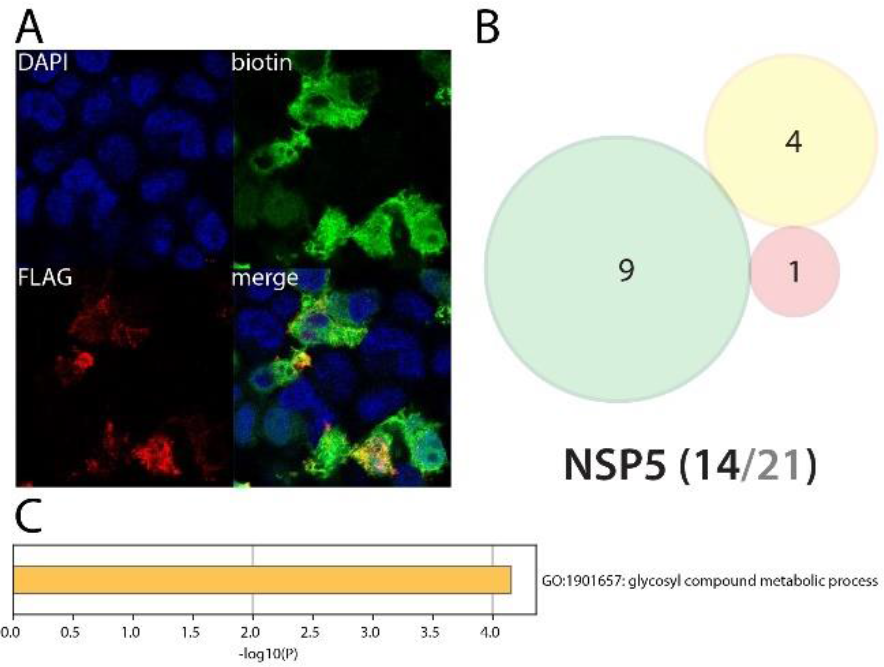
NSP5 summary. See legend Figure 1 (imaging of BirA *Flag-NSP5).

### NSP6 is a versatile membrane-associated potentially regulating lipid metabolism, autophagy and innate immune signaling

SARS-CoV-1 NSP6 has been associated with autophagosome size limitation^35^. Previous IP-MS interactomics studies of SARS-CoV-2 NSP6 show little overlap with our BioID analysis^1,2,3^. This is mainly due to the membrane-associated feature of NSP6, rendering it poorly suitable for identifying coimmunoprecipitated complexes. However, we did detect eleven previously reported interactors, including two components of the GPI-anchor transamidase complex (GPAA1 and PIGS). NSP6 proximal interactome is greatly enriched in membrane proteins (201/251), mostly from ER, mitochondria, Golgi and vesicles. Our data also uncover *e.g*. peroxisome core components (SLC25A17, PEX5, PEX11B and PEX14) and several autophagy-related proteins, including SQSTM1, which supports previous findings (by homology^35^). We detected 11 components of the NADH oxidoreductase complex, BAX, DIABLO, MFF, potentially involving NSP6 in mitochondrial respiration, apoptosis and mitochondrial fission/fusion. Our experiments also allowed the detection of several components linked to immunity response: UNC93B1, TOLLIP, ZC3HAV1, TAX1BP1, CYLD, TMEM9B, C1QBP and NPLOC4. UNC93B1 regulates several Toll-like receptors (TLRs) addressing from the ER to the endolysosome where they recognize pathogens and activates innate and adaptative immune pathways^36^. TOLLIP (Toll-interacting protein) is involved in TLR and IL-1 signaling, and acts as an adapter between ATG8 and ubiquitinated substrates, playing a major role in clearing protein aggregates^37^. ZC3HAV1 (or ZAP, Zinc finger CCCH-type antiviral protein 1) induces the recruitment of host RNA degradation machinery to the viral RNA^38^. It also positively regulates RIG-I downstream signaling, which activates IRF3 and triggers the expression of type I IFN stimulated genes (ISGs)^39^. TAX1BP1 is also involved in TNFα, IL-1 and NFκB signaling^40^. CYLD is involved in inflammation and innate immune response through its ability of inhibiting NFκB nuclear translocation^41^. TMEM9B is a key regulator of proinflammatory cytokines production in response to TNF, IL-1β and TLR ligands^42^. The Complement component 1 Q subcomponent-binding protein (C1QBP, or gC1qR) is a multifunctional protein thought to be involved in regulation of antiviral response by inhibiting RIG-I and IFIH1-mediated signaling pathways, probably through its association with MAVS after viral infection^43^. NPLOC4 acts as a negative regulator of type I IFN production via the complex formed with VCP and UFD1, which binds to RIG-I and recruits RNF125 to promote ubiquitination and degradation of RIG-I^44^. Together, these data suggest a central and versatile role of NSP6, and potentially link this viral protein to TLRs, NFκB and IL-1 signaling, apoptosis and autophagy. Poly(I:C) treatment induces NSP6 interactions with several factors involved in autophagy (SQSTM1, FAM134A, STX17), Notch (MIB2) and antigen processing (TAP2) pathways. Being associated with multiple membranes, NSP6 is at the core of a nexus of fundamental signaling processes.

**Figure 8.**
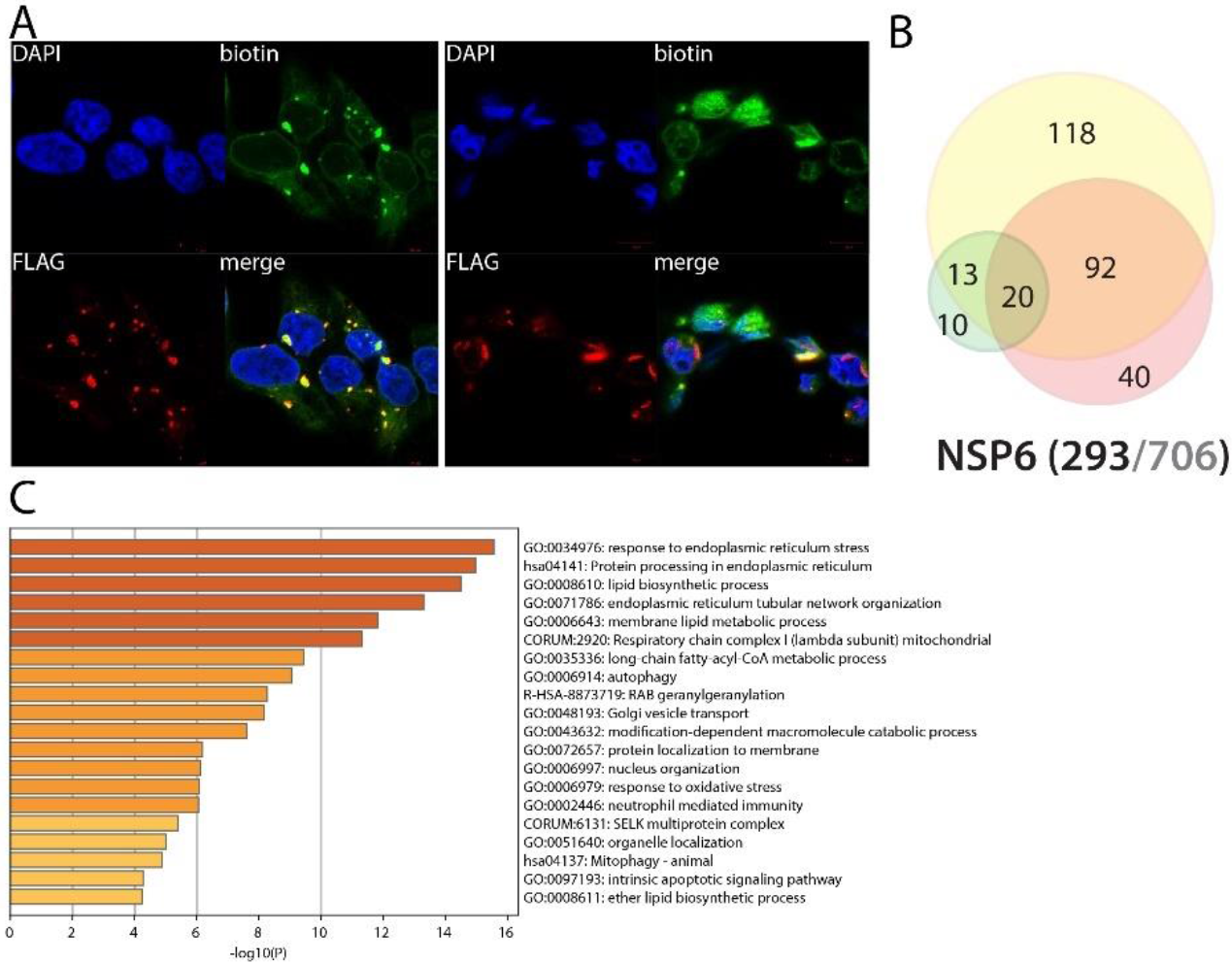
NSP6 summary. See legend Figure 1 (imaging of BirA *Flag-NSP6).

### NSP7 interacts with centrosome, microtubule cytoskeleton, vesicular trafficking machineries, nuclear factors, and could regulate TLR downstream signaling

According to SARS-CoV-1 studies, NSP7 is a subunit of the SARS-CoV-2 NSP7-NSP8 heterotetramer primase complex which is essential for an efficient NSP12 RNA-dependent RNA polymerase (RdRp) activity^45^. Hence, expressing NSP7 alone is not expected to be fully relevant, but rather indicates which factors could be recruited at the replisome through this viral protein, and potentially uncover NSP7-specific roles. The quasi-absence of overlapping between the NSP7 proximal interactome and previous NSP7 IP-MS reports (except for FOCAD and BAP1) suggests that it localizes to poorly soluble compartments. Indeed, our interactome analysis shows a strong association of NSP7 with 67 microtubule cytoskeleton associated proteins (41 centrosomal proteins), including the eight components of the HAUS complex. The presence of biotinylated vacuolar protein sorting (VPSs), kinesins (KIF16B/20A, KLC2/4) and dynamins (DNM2, DNMBP) strongly suggests that NSP7 is involved in hijacking cell vesicular trafficking. Interestingly, whereas both N- and C-ter tagged NSP7 detected interactions with cytoskeletal components, only the C-ter tagged form labeled nucleoplasmic proteins (N-terminally tagged form showed in **Fig. 9**). This observation suggests that a non-canonical nuclear localization sequence (NLS) could be localized in the N-ter region of NSP7 (impaired by the N-ter tag), or that NSP7 can be cleaved, with a C-ter fragment translocated in the nucleus. This discrepancy is quite uncommon for a nonmembrane associated protein, and further experiments are required to elucidate the molecular etiology. NSP7 nuclear interactors were essentially involved in DNA-templated transcription, DNA repair, mRNA processing and histone modification. These results suggest that NSP7 could impair host mRNA production, thus precluding proper antiviral transcriptional program and/or reducing the pool of endogenous mRNA (*i.e*. lighten host mRNA translational burden and enhance host cell capacity for viral RNAs translation). The C-ter tagged NSP7 also specifically captured important innate immune and inflammatory responses modulators under basal culture conditions, such as HERC5, PPM1B, TRIM56 and TBK1. Of interest, TBK1 (Serine/threonine-protein kinase TBK1) is activated by TLR following viral or bacterial sensing. TBK1 then associates with TANK and TRAF3 and phosphorylates the IFN regulatory factors 3 and 7 (IRF3 and IRF7). These modifications lead to the activation of pro-inflammatory and antiviral genes, including type I IFN^46–48^. PPM1B is a phosphatase able to dephosphorylate TBK1, inhibiting its activation^49^. The pre-treatment with poly(I:C) did not induce major NSP7 interactome rewiring.

**Figure 9.**
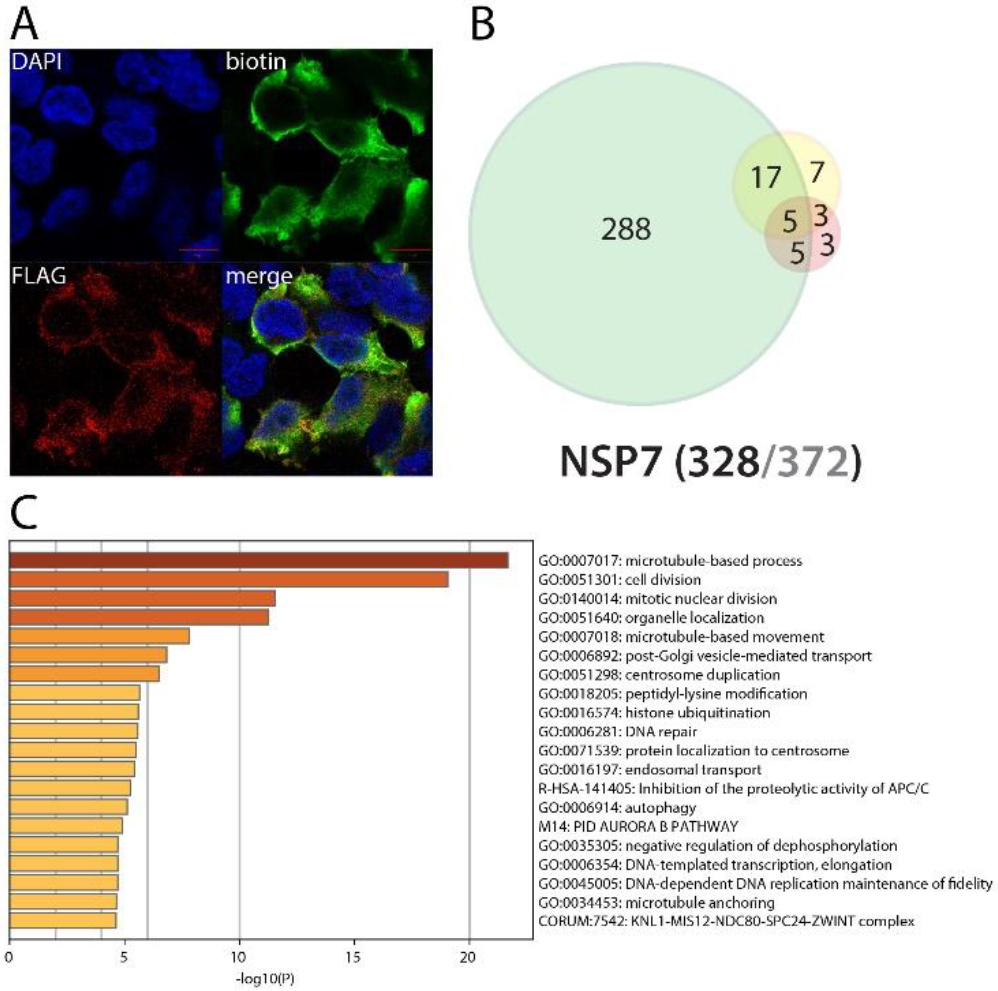
NSP7 summary. See legend Figure 1 (imaging of BirA *Flag-NSP7).

### NSP8 partners with HECT-domain containing proteins and possibly with DNA in the nucleus

Together with NSP7, it forms the viral primase complex. Unlike NSP7, NSP8 did not detect many proximity interactors, suggesting that it either functions only in actual infectious context or with its viral partners, and could be dragged to proper locations through the NSP7 subunits. The only noticeable interactors of NSP8 are the E3 ligases HECTD1 and HERC1 (both reported^2^). C-ter tagged NSP8 interacts with PSMC3IP and histones (H2A6, H1-0 and H1-2) which might indicate an association with DNA that is in line with the observed localization of NSP8 interactors in the nucleus (**Fig. 10**). These interactions could suggest a preferential binding of NSP8 to HECT-domain containing protein, with a so far unknown significance. Poly(I:C) did not induce any interaction gain other than FBXO30 and MTR.

**Figure 10.**
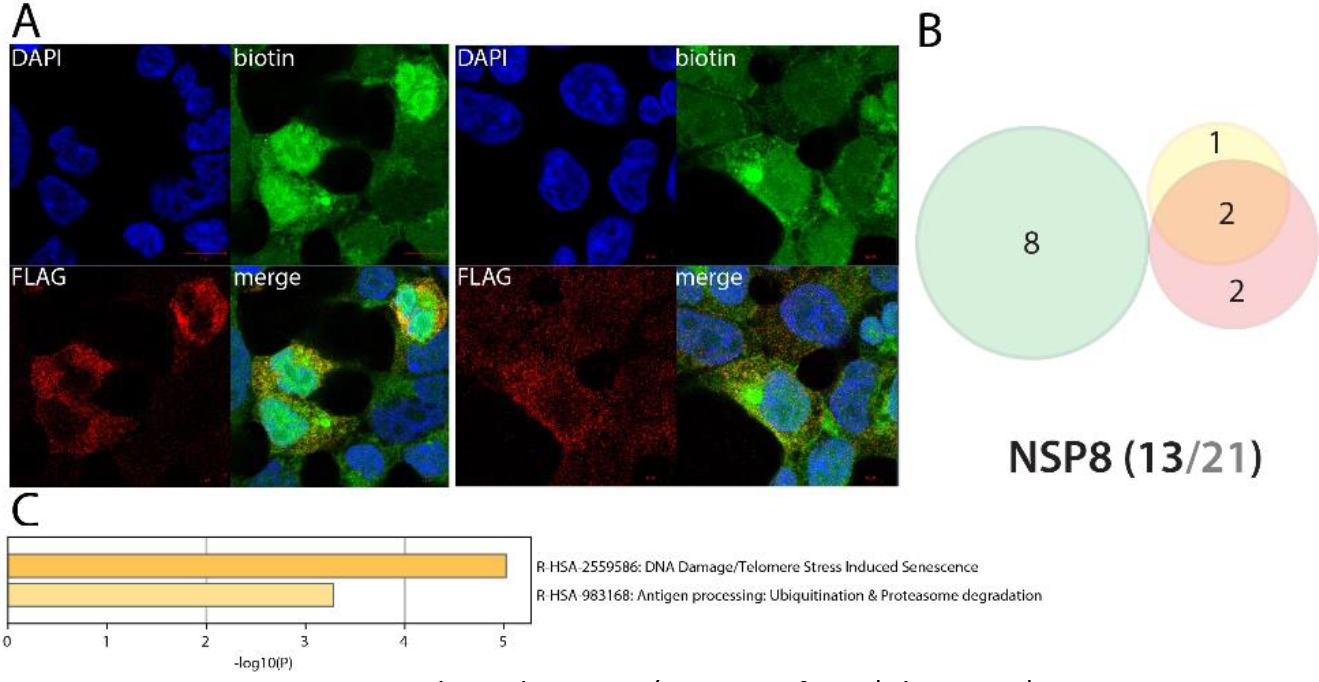
NSP8 summary. See legend Figure 1 (imaging of BirA*Flag-NSP8).

### NSP9 could impair endogenous mRNA processing and mitosis

NSP9 (RNA replicase) is predicted to mediate viral replication and virulence^50^. Only two interactors were previously reported (GTF2F2 and SETD2). Amongst the interactors identified by BioID, we detected several RNA processing factors (FMR1, FXR1 and FXR2) involved in mRNA capping and transport via the nucleopore. The presence of myosin (MYH10) and tropomyosin (TPM1) could indicate a role of NSP9 in cytoskeleton contractility. BioID data obtained in basal culture conditions did not reveal other highly specific features. Poly(I:C) treatment did not induce noteworthy changes in NSP9 interactions.

**Figure 11.**
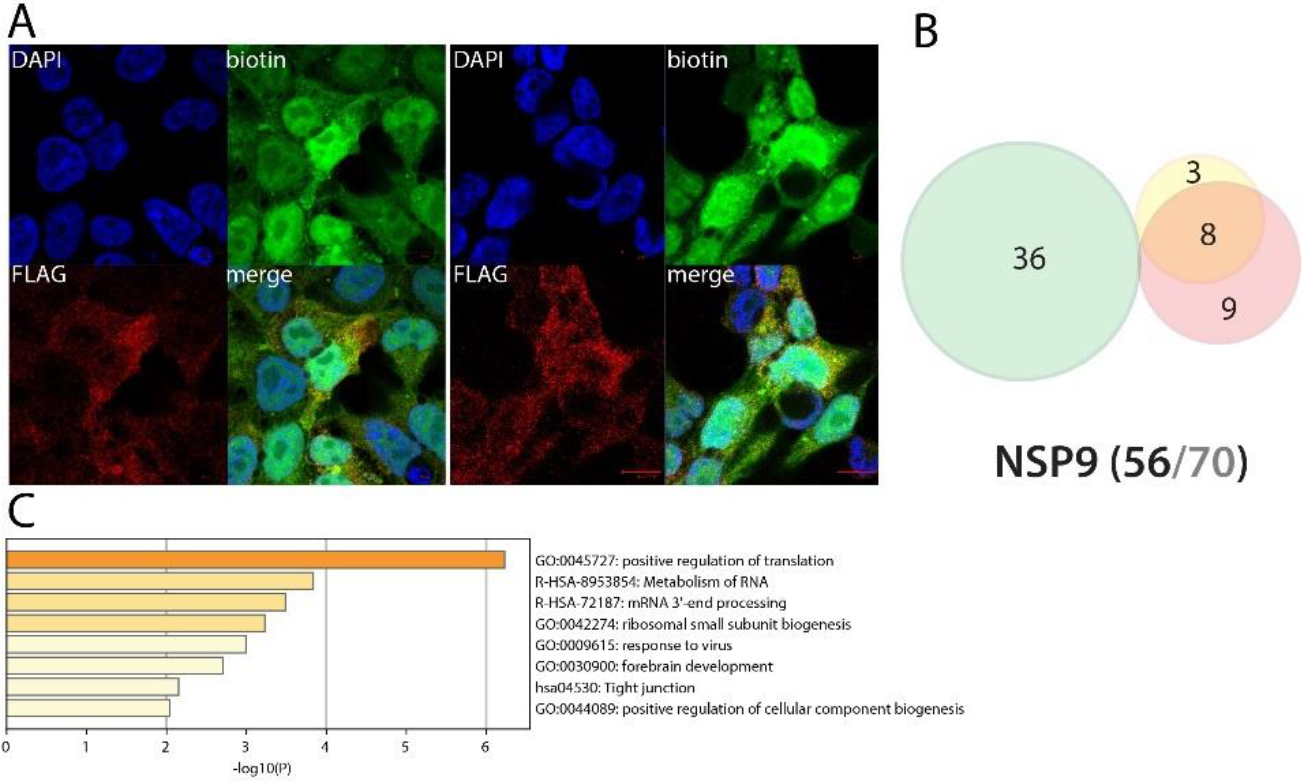
NSP9 summary. See legend Figure 1 (imaging of BirA*Flag-NSP9).

### NSP10 alone interacts with nuclear factors involved in chromatin remodeling and mRNA processing

NSP10 is a cofactor of both NSP14^51^ and NSP16^52^, stimulating their 3’-5’ exoribonuclease and 2’-O-methyltransferase activities, respectively. No previously reported NSP10 interactors were identified in our analysis. According to our immunostaining, more than half of NSP10 BioID interactors are nuclear, including chromatin remodelers, transcription factors and RNA processing proteins. Surprisingly, N-ter BioID tagged NSP10 did not localize in the nucleus. We thus hypothesize it could interfere with nuclear factors either which shuttle to the nucleus or upon mitosis. Of note, NSP10 detected TLE1, an inhibitor of NFκB regulated gene expression^53^ and AZI2 (or NAP1), another regulator of NFκB-dependent antiviral innate immune response^54^. Alike NSP7-NSP8, NSP10 expression in the absence of its viral partners NSP14 or NSP16 renders this interactome hard to interpret. Poly(I:C) treatment did not induce noticeable gains of NSP10 interactors.

**Figure 12.**
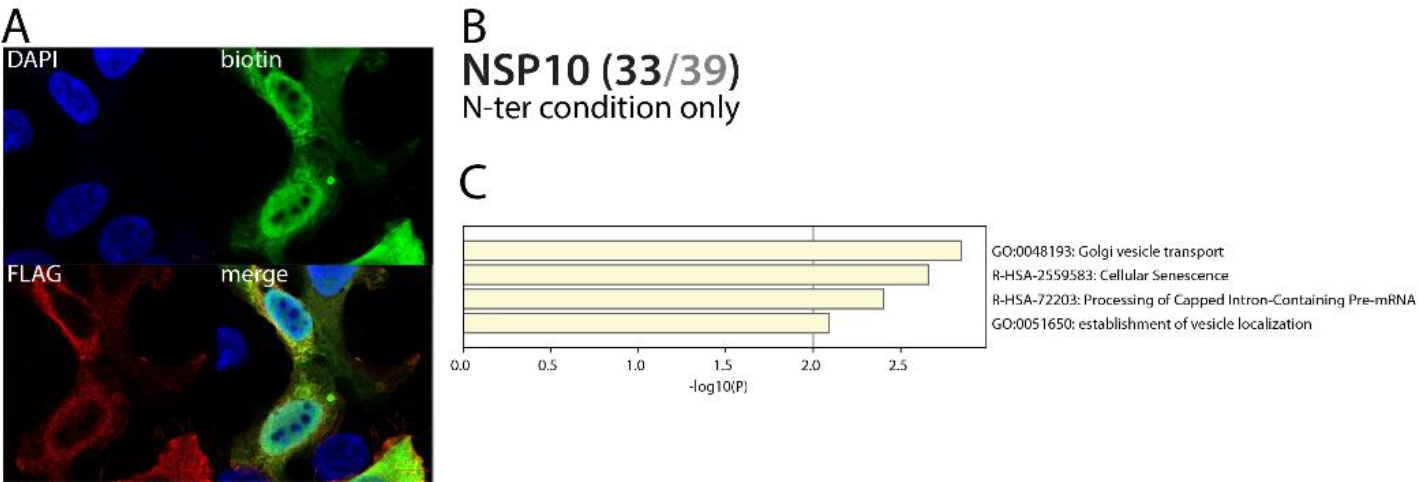
NSP10 summary. See legend Figure 1 (imaging of BirA*Flag-NSP10).

### NSP12 (RdRp) associates with RNA polymerase cofactors, but also with antiviral and inflammatory signaling proteins

NSP12 is an RdRp binding NSP7-NSP8^55^. Due to its crucial function in viral RNA replication^56^, NSP12 is a target of choice for antiviral therapeutic strategies. Tremendous efforts have thus been deployed to target this RNA polymerase, and several clinical assays (*e.g*. Remdesivir^57^) are in process to reposition antiviral drugs and combat COVID-19 severity. It is thus of the utmost importance to finely characterize its interactors to provide further knowledge in how this viral protein behaves in host cells. Our BioID experiments identified microtubule-associated proteins (centrosomal, centriolar satellite and cilia), RNA-mediated gene silencing machinery (AGO2, TNRC6A, TNRC6B), chaperones (*e.g*. CCT2/5/6A/8, FKBP4, DNAJC7, HSP90A1/A1B/A8, VBP1, PFDN1/2/4/5/6), anaphase-promoting complex components (ANAPC1/4/5/7/13/16, CDC16/23/27), phosphatidylinositol regulating proteins (PIK3R2, PIKFYVE, INPPL1) and JNK/MAPK signaling pathway regulators (*e.g*. MAP3K7, PEAK1). In addition to these enriched categories, several proteins associated with RNA polymerases were identified: POLR2E/L, RPAP3, INTS4/7, PIH1D1, ENDOG and URI1, CPSF3, CSTF2T. They represent either candidate cofactors recruited to enhance NSP12 activity or proteins targeted to impair host transcription. Several proximal interactors are also likely to link NSP12 to innate immune response modulation: TRIM56, TRAF3IP1, CYLD, RIPK1, TAB1, TAB2, and TBK1. TRAF3IP1 inhibits IL-13 signaling by binding to IL13RA1, which is involved in allergic inflammation and diseases such as asthma^58^. The multiple symptoms triggered by IL-13 signaling (airway hyperresponsiveness, mucus hypersecretion) could resemble the acute respiratory distress frequently occurring in severe COVID-19. TAB1 mediates signaling between TGFβ receptor and MAP3K7, and TAB2 acts as an adapter between MAP3K7 and TRAF6 and promotes the activation of MAP3K7 downstream IL-1 activation^59,60^. Besides the gain of the IFN-induced protein with tetratricopeptide repeats 2 (IFIT2), no obvious NSP12 interactome change was observed following poly(I:C) treatment. Together, these interactors could link NSP12 to a wider range of signaling processes than expected, including antiviral and inflammatory pathways.

**Figure 13.**
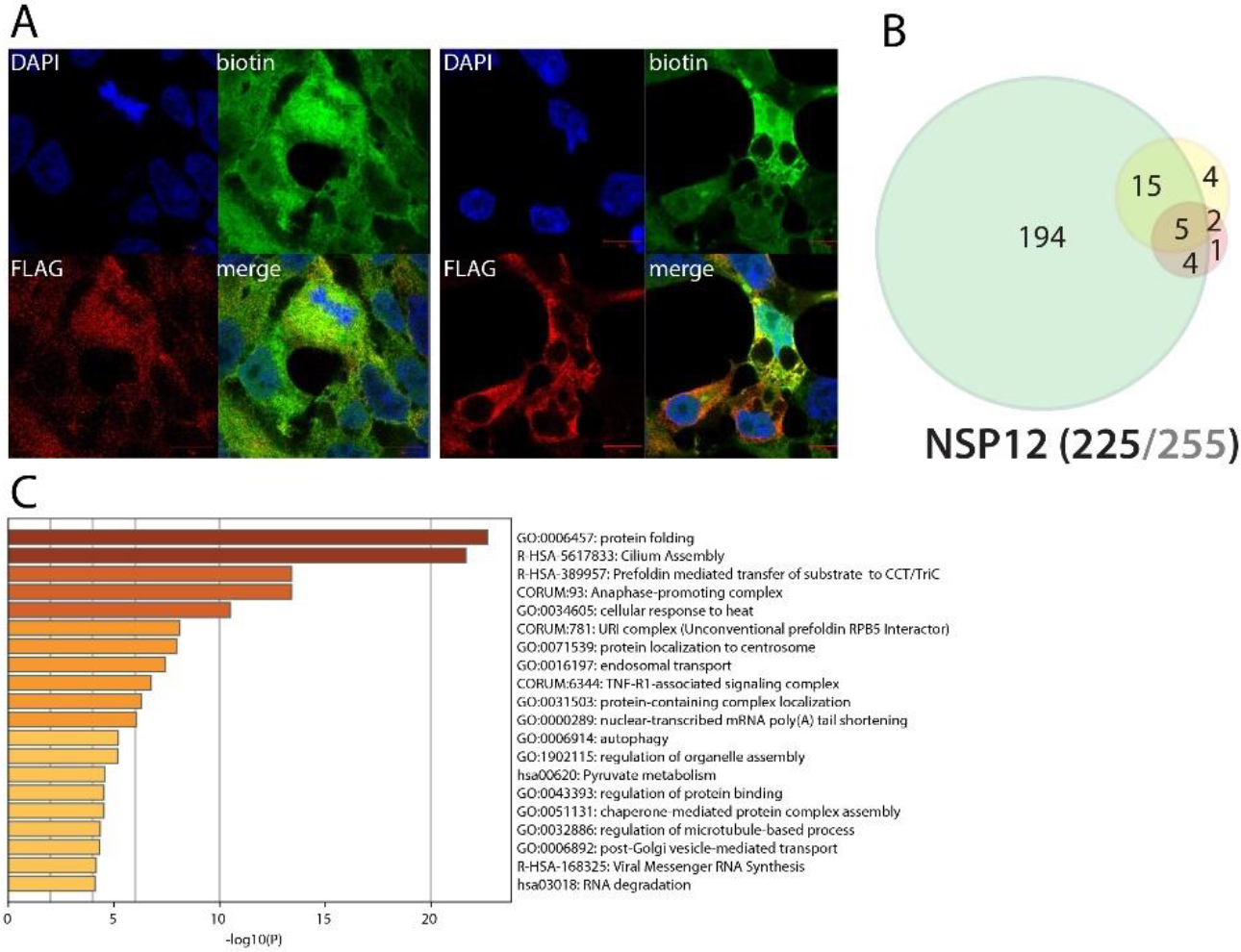
NSP12 summary. See legend Figure 1 (imaging of BirA *Flag-NSP12).

### NSP13 (helicase) alone associates with centrosomal components

The NSP13 helicase unwinds the SARS-CoV-2 dsRNA (which transiently forms upon vRNA replication) into two ssRNA to make it suitable for RNA replication and displays a dsDNA helicase activity^61^. SARS-CoV-1 NSP13 is also able to hydrolyze deoxyribonucleotide and ribonucleotide triphosphates^62^. Previous interactomics data showed an association between NSP13 and centrosomal proteins. Although we did not identify the same partners, our proximal interactome was also enriched in centrosome components (*e.g*. CCP110, CEP76, POC1B). Besides these interactions, we did not detect specific features revealed by proximal partners enrichment analysis, regardless of the poly(I:C) treatment. Importantly, a recent functional study has suggested that SARS-CoV-2 NSP13 is an IFN signaling antagonist^63^. However, we did not detect any NSP13 interactor which could explain this phenotype. We hypothesize that this role can be dependent on the molecular context. Future experiments in SARS-CoV-2 infected cells will reveal additional and pathologically relevant interactions of this essential viral protein.

**Figure 14.**
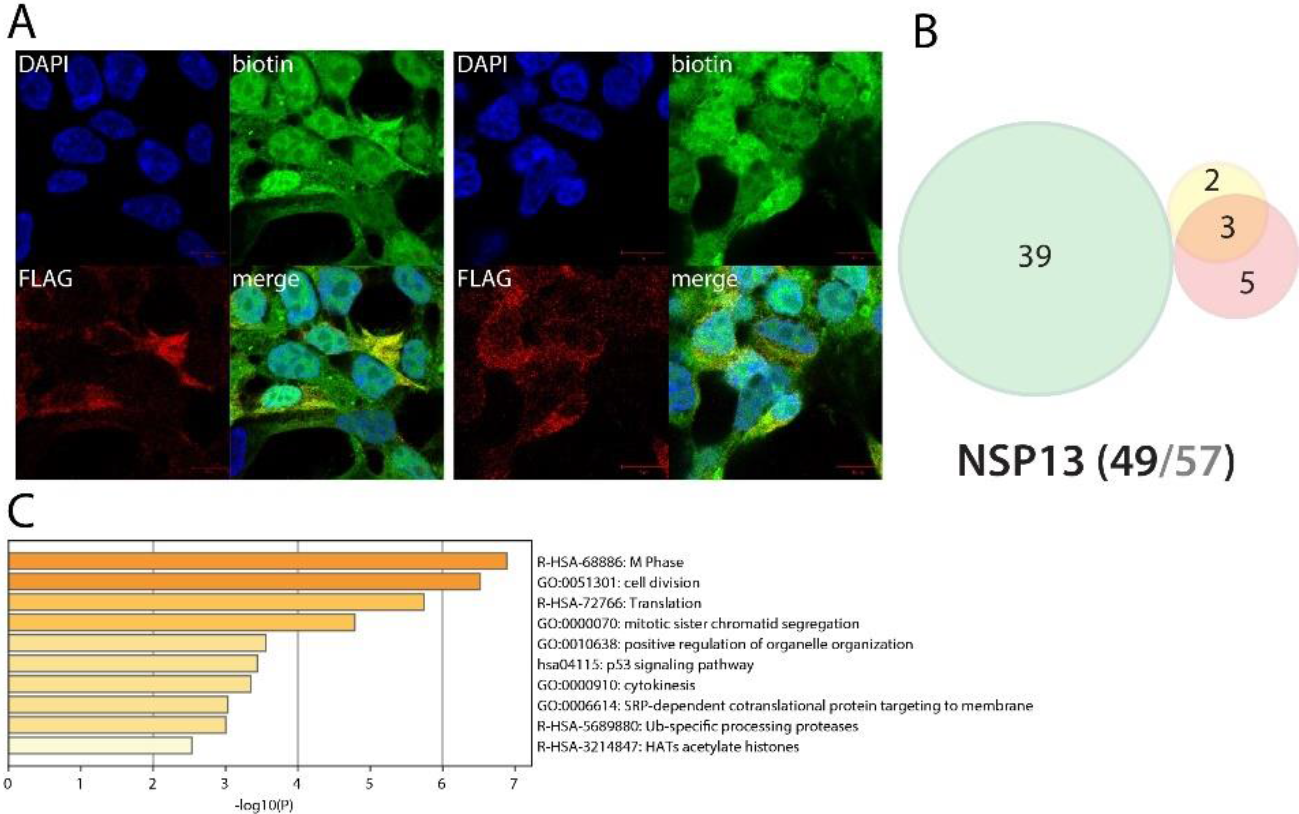
NSP13 summary. See legend Figure 1 (imaging of BirA *Flag-NSP13).

### NSP14 associates with RNA decapping and deadenylation host factors

NSP14 is a component of the replication-transcription complex. It is a dual functional enzyme, bearing both 3’-5’ exonuclease (ExoN) and a guanine N7-methyltransferase (N7-MTase) activities^64^. ExoN ensures a proofreading function responsible of high-fidelity SARS-CoV-1 replication through its ssRNA and dsRNA hydrolyzation activity. The N7-MTase function is required for vRNA capping, precluding the triggering of host non-self RNA detection system^64^. Only SIRT5 is common between our proximal interactomic study and previous reports^2^. Our proximal interactome reveals several interactions with RNA-associated proteins: DCP1B, MEX3A, PATL1, CNOT2 and PAN2. Through these partners, NSP14 could interfere with RNA decapping and deadenylation, two processes leading to mRNA degradation. Further experiments are needed to get precise insights on how NSP14 functions, particularly performing interactomic studies in infected cells, or at least in cells expressing its viral cofactor NSP10. NSP14 uniquely interacts with TRAF7, an E3 ubiquitin ligase involved in NFκB activation^65^. Upon poly(I:C) stimulation, we detected an interaction between NSP14 and CEP97/152/192, three proteins involved in ciliary basal body docking to the plasma membrane. The enriched interactions with machineries leading to mRNA degradation could explain why NSP14 was also identified as an IFN response antagonist^63^. We could hypothesize that NSP14 could either sequester host decapping and deadenylation machineries or stimulate host mRNA degradation.

**Figure 15.**
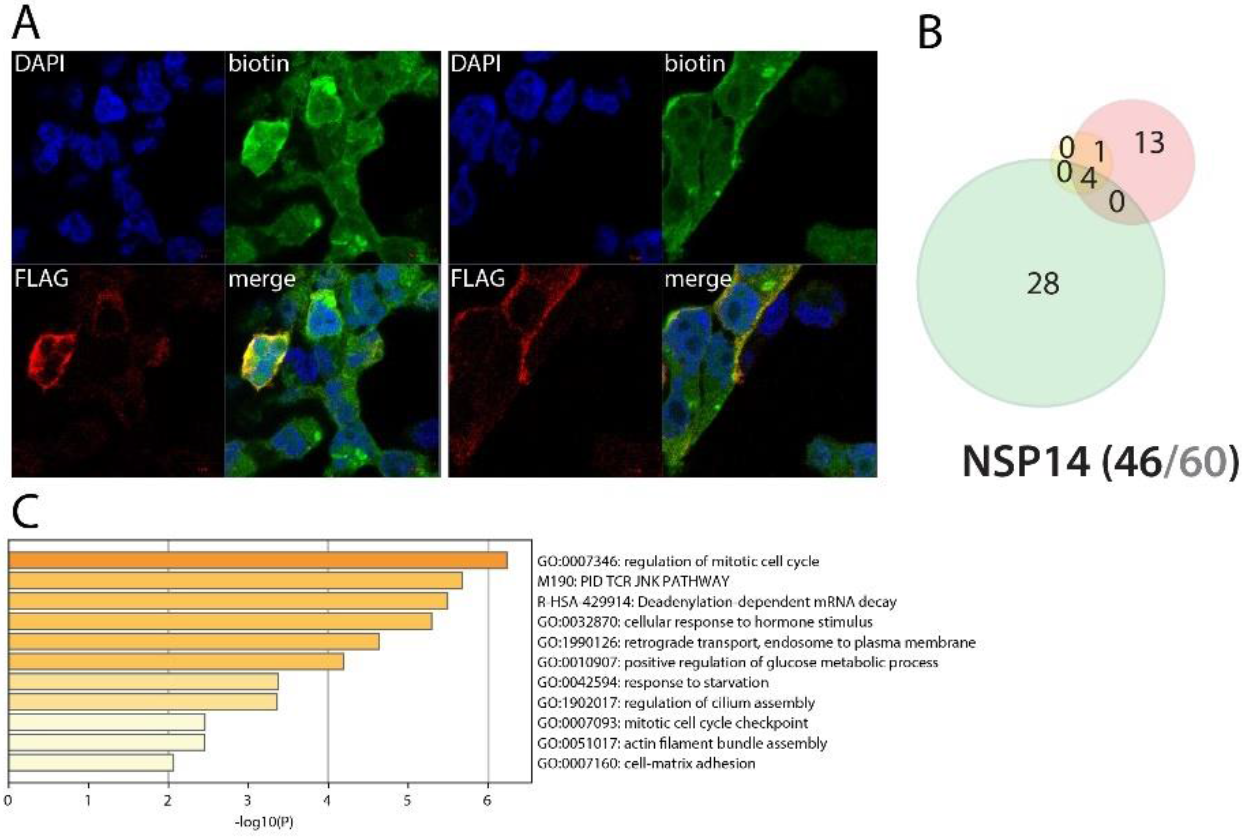
NSP14 summary. See legend Figure 1 (imaging of BirA *Flag-NSP14).

### NSP15 interacts with innate immune response components downstream TLRs and might have an impact on mRNA decay and apoptosis regulation

NSP15 is an RNA uridylate-specific endoribonuclease with a C-terminal catalytic domain from the EndoU family^66^. It trims ss/dsRNA, producing 2’-3’ cyclic phosphodiester and 5’-hydroxyl termini. Recent studies suggest that NSP15 may interfere with the innate immune response, possibly antagonizing dsRNA sensing in macrophages^63^. There is no overlap between our NSP15 BioID studies and previous interactomic works^1,2,3^. NSP15 BioID identified several physical and functional entities, *e.g*. prefoldin complex (PFDN1/2/4/5/6, VBP1), nonsense mediated mRNA decay factors (SMG5/6/7/9), and actin cytoskeleton associated components. Since NSP15 has also been reported as impairing adequate innate immune response, we focused our analysis on host-factors linked to these mechanisms. Our dataset reveals novel interactions between NSP15 and RIPK1, TAB1, TAB2, TRAF3IP2 (CIKS) and XIAP. These proteins are activated downstream TLR activation and/or are involved in the IL-17 signaling pathway. Across our dataset, DIABLO and HTRA2 were identified as NSP15 high confidence proximal interactors, potentially linking NSP15 to apoptosis regulation. Beyond its EndoU reported function, our data strongly suggest that NSP15 impacts host cell innate immune signaling and apoptosis, interfering with signal transducers downstream TLRs and DIABLO/XIAP/HTRA2, respectively. Under poly(I:C) stimulation, NSP15 gained a handful of interactors, but none with infection-related function. Together, our data shed a new light on how NSP15 could interfere with host innate immune response and suggest its involvement in apoptotic pathway deregulations.

**Figure 16.**
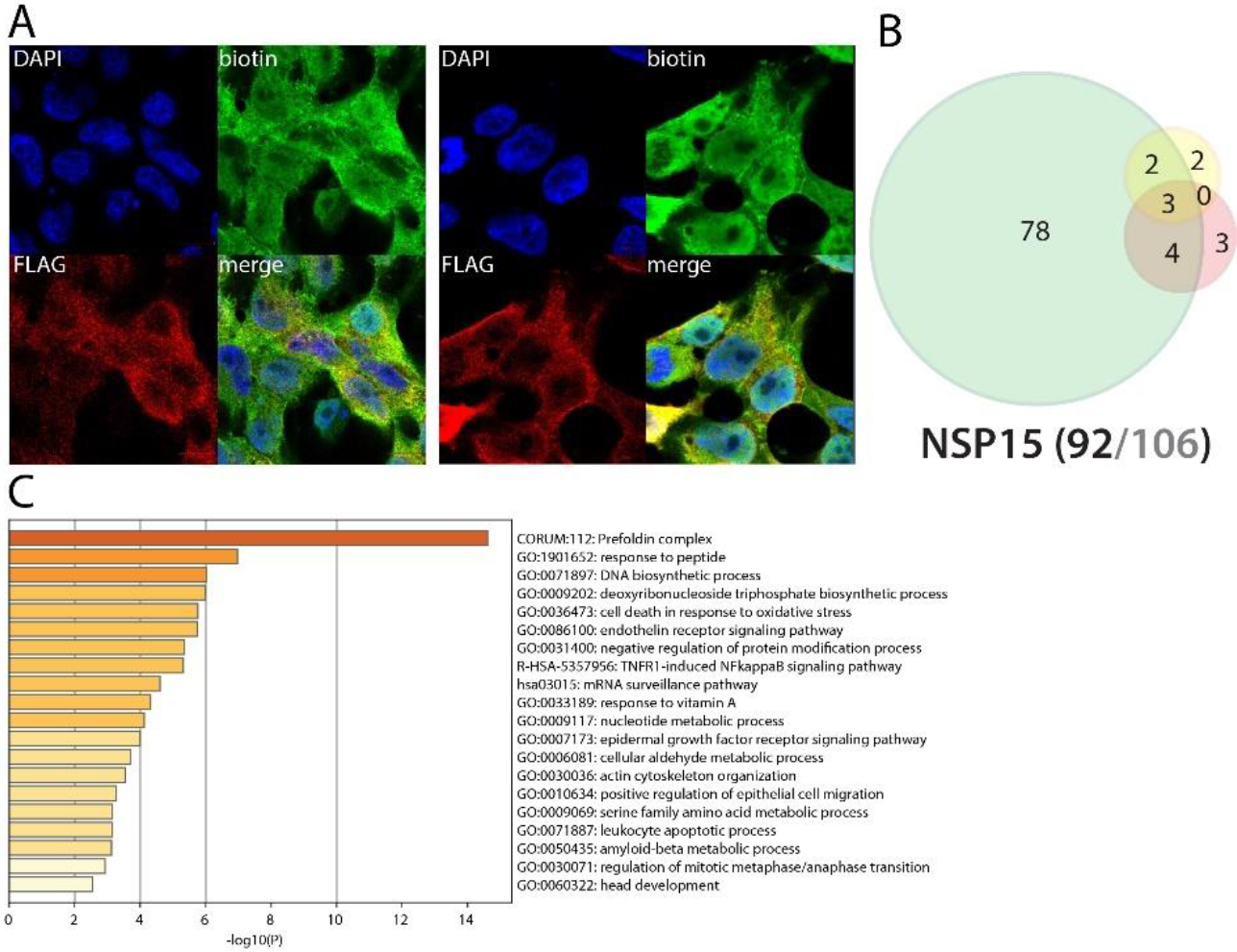
NSP15 summary. See legend Figure 1 (imaging of BirA*Flag-NSP15).

### NSP16 interacts with multiple components controlling NFκB, IFN, CTNNB1 and CREB nuclear translocation, potentially impairing proper cell response to multiple signals

NSP16 protein acts as a s-adenosylmethionine-dependent (nucleoside-2’-O)-methyltransferase in the presence of its partner NSP10^67^. NSP16 is involved in several biological processes such as viral signal transduction, nucleic acid processing, chromatin remodeling, metabolism, detoxification, and mRNA capping. No overlap was observed between our study and previous reports^1,2,3^. NSP16 proximal interactors analysis showed an enrichment in components of the: nuclear pore complex; APC/C, centriole; kinesin complex (KIF7/13A/13B/14/20A, KLC4); nucleus and P-body RNA-processing machineries (AGO2, DCP1A, XRN1, PATL1, TNRC6B, MEXA3, PAN2, EIF4ENIF1). DCP1A and PATL1 are interesting candidates to link NSP16 to mRNA capping process. In addition, NSP16 uniquely identify the p100 NFκB subunit (NFKB1), which could lead to defect in NFκB signaling or transport into the nucleus. Together with the detected nucleopore components, NSP16 interactions with NFκB factor, TBK1, CRTC1 (regulating CREB transcription factor^68^), b-Catenin (CTNNB1) and CTNNB1 regulators APC^69^, nuclear pore components (NUP37/98/107/133/L1) strongly suggest that NSP16 could generally interfere with the nuclear translocation of transcription factors following activation of several activating pathways. Poly(I:C) stimulation induces interactions with DTX3L-PARP9, additionally linking NSP16 to IFN response regulation^20^. Our data thus open numerous hypothesis on additional roles of NSP16 towards several signaling pathways, including antiviral response.

**Figure 17.**
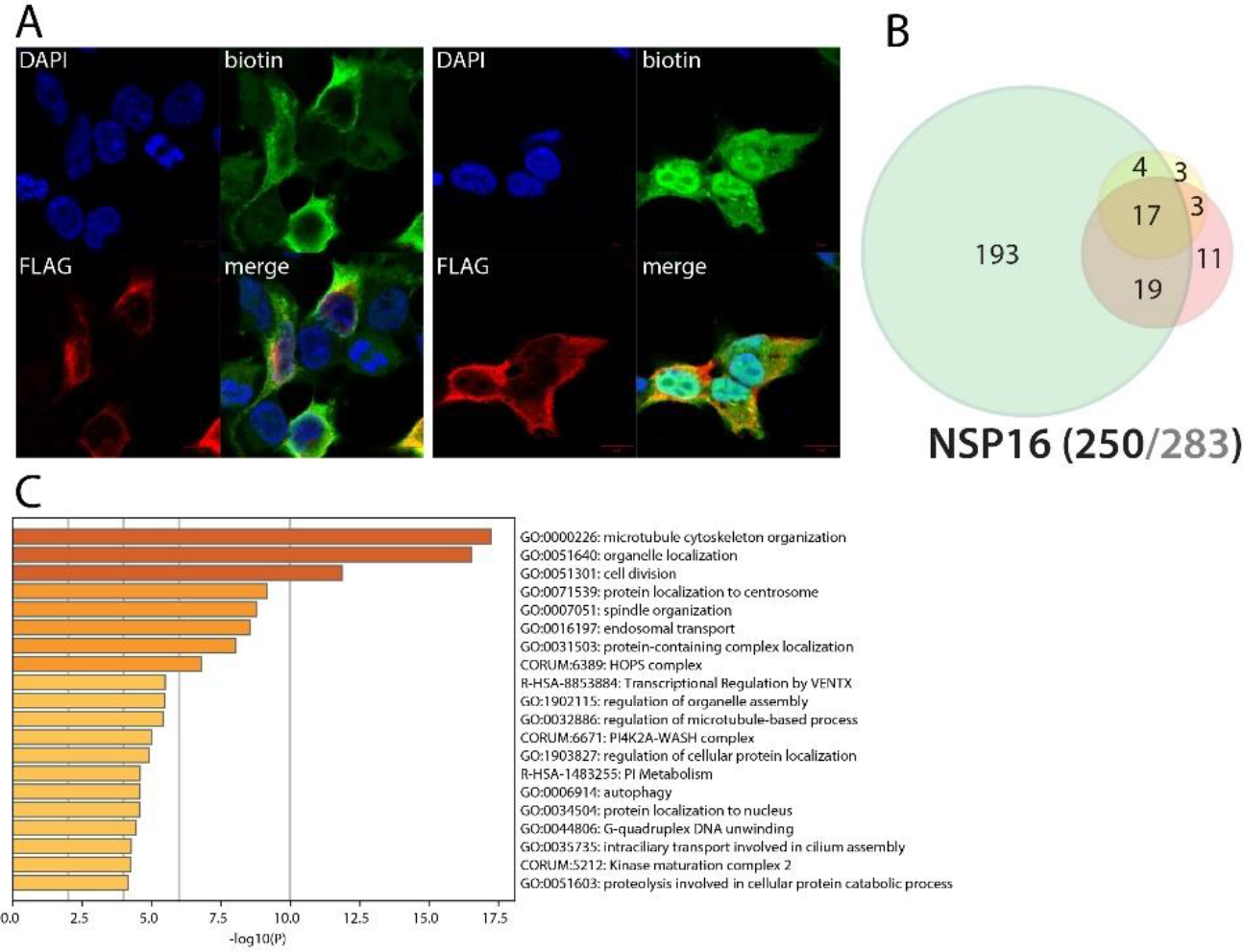
NSP16 summary. See legend Figure 1 (imaging of BirA*Flag-NSP16).

### Spike (S) intracellular interactors are involved in cell adhesion and migration

The Spike glycoprotein (S) is responsible of virion attachment to host cell through interacting with ACE2 and TMPRSS2^70^. S proteolysis by cathepsin unmasks the fusion peptide S2 and activates membrane fusion with endosomes^71^. Despite tremendous efforts to understand and target this SARS-CoV-2 fundamental structural protein, S functions inside cells remain largely unknown. It is reported that S can accumulate at the ER-Golgi intermediate compartment, where it participates in virus assembly^72^. Eventually, S oligomers might be transported to the plasma membrane and promote cell-cell fusion^73^. Expectedly, we identified S proximal interactors mostly assigned to ER, Golgi, endosome and plasma membranes. The C-ter tagged version of S provided the richest results, suggesting that the N-ter tag impairs proper S addressing (probably through masking the signal peptide or because of the BioID-tag cleavage from the fusion). We identified two previously reported interactors of S: EZR and ZDHHC5. Of note, we identified 12 factors involved in virus entry mechanisms. We detected highly specific interaction of S with four components of the Ragulator complex (LAMTOR1/2/3/5). S BioID detected PPP1R16B, CSPG4, and JUP, which are all involved in endothelial cell adherence and motility. Additionally, S is in proximity of multiple factors involved in cell-cell and cell-matrix adhesion (*e.g*. JAM3, CASK, EPB41L4B, FLRT3, FLOT2, CD44, TENM3), lamellipodium (*e.g*. ROCK1, RDX, PKN2) and migration (MARCKS, ROBO1/2, DAG1). This set of interaction could be related to the recently reported SARS-CoV-2-driven induction of cellular extension/pseudopodia, which are thought to facilitate cell-to-cell virus spreading^18^. Finally, S interacts with several factors involved in inflammation and innate immunity regulation, such as ITCH, TOLLIP, HGS and a set of kinases involved in NFκB activation (PRKCI^74^, PRKD1^75^, PRKCQ). Poly(I:C) pre-stimulation did not induce any noticeable interaction.

**Figure 18.**
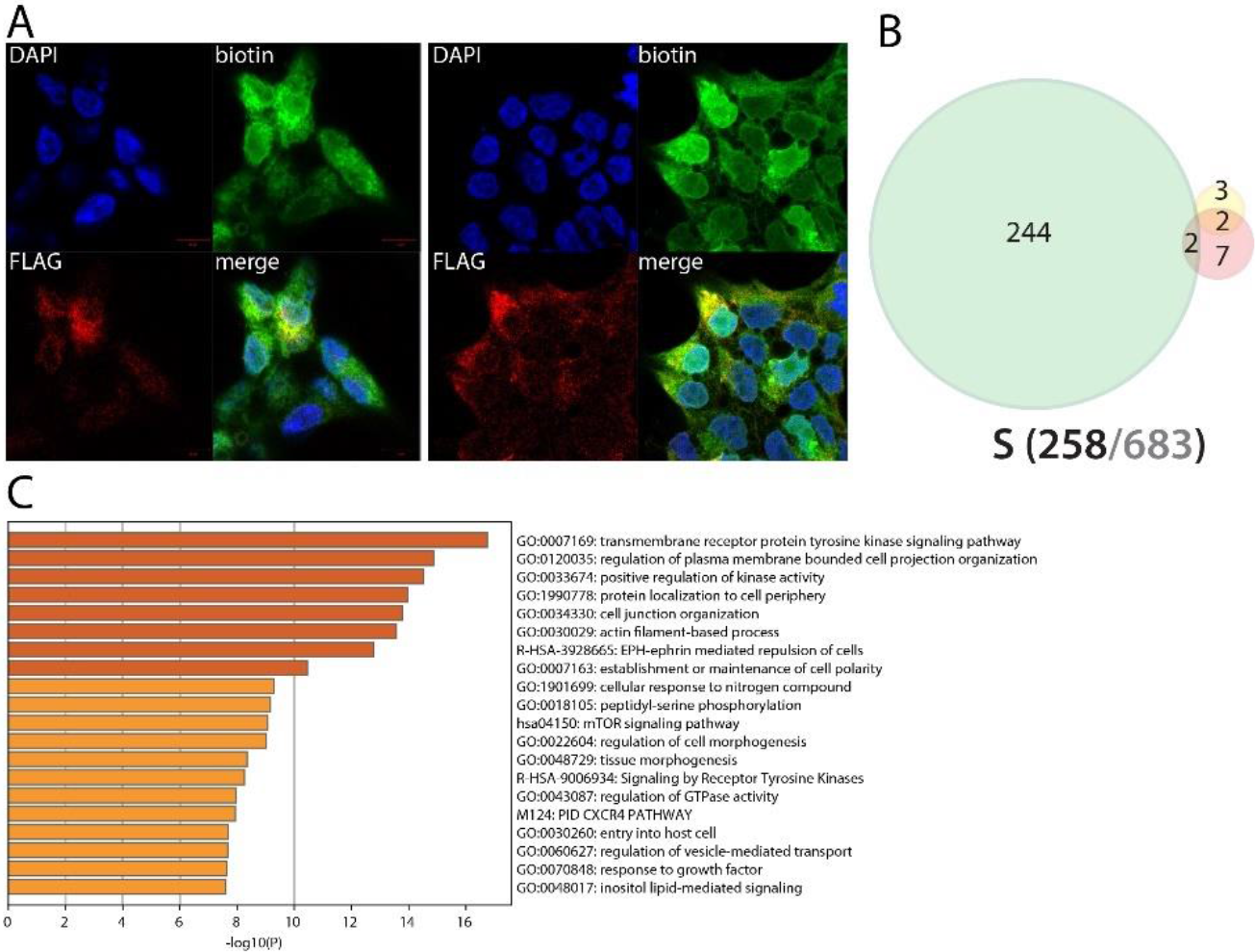
Spike (S) summary. See legend Figure 1 (imaging of BirA*Flag-S).

### ORF3a interacts with vesicular sorting and fusion machineries, and could impact a plethora of membrane proteins, such as receptors involved in thrombosis, type II IFN, NFκB and IL-6 signaling

ORF3a is a key accessory protein showing a pro-apoptotic activity^76^. By homology with SARS-CoV-1, ORF3a is a Golgi transmembrane protein displaying a Na+ or Ca2+ ion channel activity and is implicated in NLRP3 inflammasome activation through TRAF3-dependent ubiquitination of ASC (the adapter protein of the inflammasome)^77^. While non-essential for SARS-CoV-1 infection, it plays a role in virulence^78^. We identified 11 previously reported SARS-CoV-1/2 interactors. Our N-terminally tagged ORF3a BioID dataset is enriched in ER stress response and misfolded protein pathway, probably reflecting a processing impairment due to the tag positioning. Over 80% of ORF3a interactors are related to membranes, mostly Golgi, plasma membrane and vesicles. ORF3a detects 7 components of the SNARE complex (NAPA, STX4/6/7/12/16 and SNAP23) and uniquely interacts with 12 components of the TRAPP complex (TRAPPC1/2/2L/3/4/5/6B/8/9/10/11/12). Several CORVET and HOPS complex components were also identified (VPS8/16/18/33A/39/41). The SNARE complex is involved in membrane trafficking and fusion^79^; the TRAPP complex in tethering vesicle transported from ER to Golgi^80^; and the HOPS and CORVET complexes in vacuolar, late endosomal and lysosomal membranes tethering and fusion^81^. These data strongly suggest that ORF3a acts as a major regulator of endosomal and membrane trafficking by interfering with vesicular membrane attachment machineries. Through interacting with endosomal addressing machinery, ORF3a potentially impacts the fate of a plethora of membranes and their embedded proteins. Among interactors potentially related to reported ORF3a functions and COVID-19 symptoms, we identified: NUMBL, TNFRSF10B, SCRIB, UNC5B, SRC, ADAM9, IL6ST, TOLLIP, ROR1, and PTPRJ. The Numb-like protein (NUMBL) inhibits the NFκB pathway, inducing TRAF6 degradation^82^. TNFRSF10B (Tumor necrosis factor receptor superfamily member 10B, or CD262) is the TRAIL ligand receptor and activates the Cas8-dependent apoptotic pathway^83^. It has also been reported to promote NFκB pathway activation. SCRIB^84^ and UNC5B^85^ can be linked to polarity and apoptosis regulation. SRC (Proto-oncogene tyrosine-protein kinase Src) is a core signaling protein downstream immune response,adhesion and cytokine receptors^86^. Of note, we detected other related kinases, such as LYN and YES1. LYN is involved in innate and adaptative immune responses, transducing signal downstream *e.g*. TLR2/4/6, cytokine receptors (CXCR4, receptors for IL-3, IL-5, CSF2) and growth factors in platelets, neutrophils and eosinophils (*e.g*.^87,88^). ADAM9 (Disintegrin and metalloproteinase domain-containing protein 9) cleaves multiple substrates involved in angiogenesis, may act as an alpha secretase for APP and has been involved in viral infection mechanisms^89^. IL6ST (Interleukin-6 receptor subunit beta) is involved in transducing multiple signals, such as IL-6 and IL-11, and might be crucial in severe COVID-19 cases^90^. ROR1 (Inactive tyrosine-protein kinase transmembrane receptor ROR1) acts as a receptor, and when activated by WNT5A ligand binding, activates NFκB signaling^91^. Finally, PTPRJ (Receptor-type tyrosine-protein phosphatase eta; CD148) regulates endothelial cell survival, enhancing epithelial cell junctions as well as VEGF-induced SRC and AKT activation^92^. PTRJ has also been reported as activating platelets and triggering thrombosis (as well as F11R)^93^. These interactors suggest a central role of ORF3a in dysregulating membrane receptor-containing endosomes, which could lead to the alteration of multiple pathways such as apoptosis signaling, innate and adaptative immune responses and platelet activation. No major interactors gain was observed in the presence of poly(I:C).

**Figure 19.**
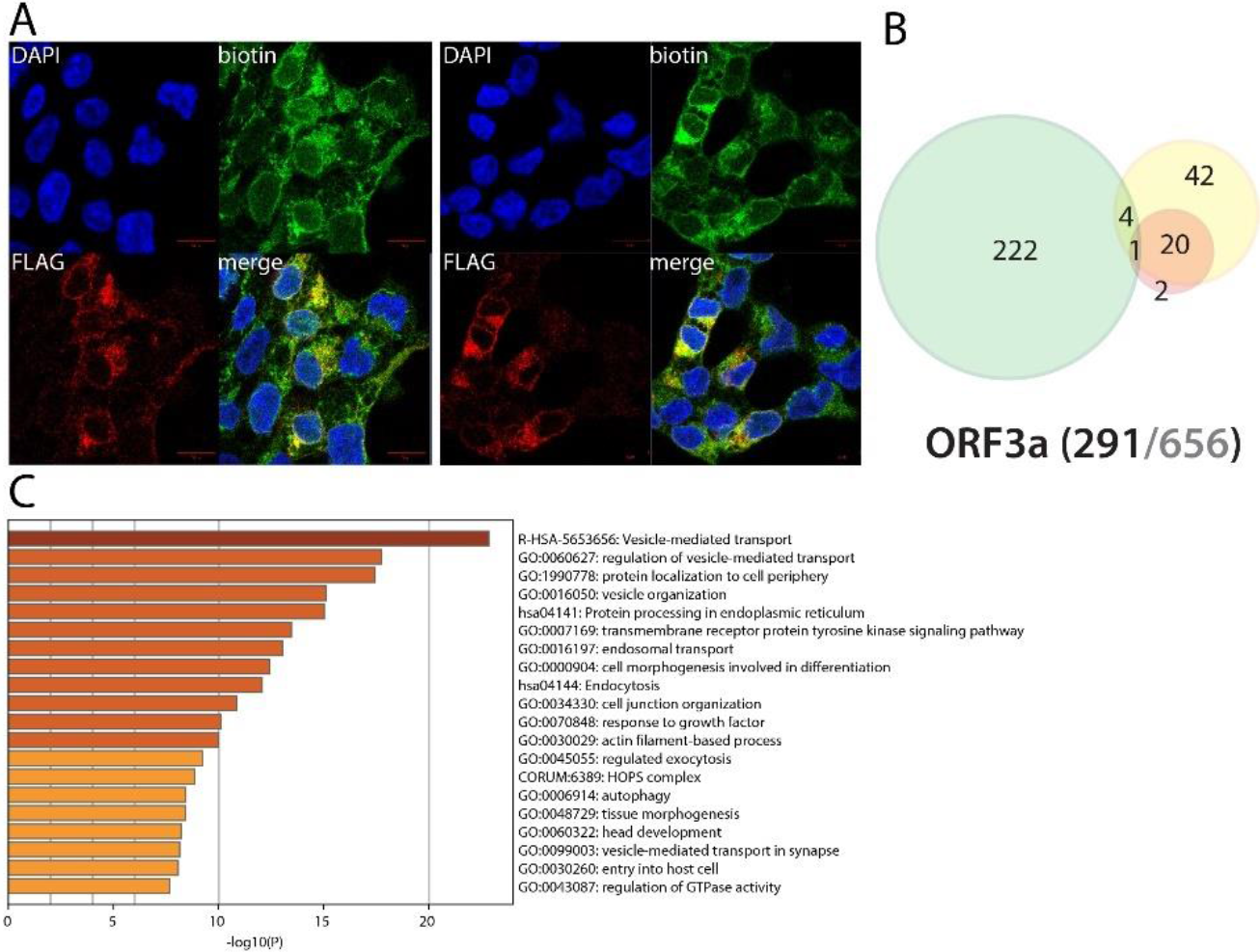
ORF3α summary. See legend Figure 1 (imaging of BirA *Flag-ORF3a).

### ORF3b is a Golgi-associated protein interacting with ESCRT-0

Little is known on ORF3b function. Unlike SARS-CoV-1 ORF3b, SARS-CoV-2 ORF3b shows type I IFN antagonist potential^94^. Comparing with previous reports, we identified four interactors in common: STAM2, EPHA2, STX6 and PTPRJ. Alike ORF3a, ORF3b is a Golgi resident protein (47/150 hits) interacting with multiple vesicle-related proteins and membrane components. ORF3b could thus also been involved in membrane receptor fate impairment, and thus target numerous pathways. Our BioID experiments identified the three components of the ESCRT-0 complex: HGS, STAM and STAM2, a vesicle sorting complex which directs endosomes towards lysosome and is crucial in multivesicular body formation. These proteins are dedicated to downregulating membrane receptors abundance^95^. Interestingly, the ESCRT-0 complex has also been involved in IL-2 and GM-CSF/L signal transduction^96^. ORF3b also identified multiple components of the SNARE complex, but did not detect HOPS, CORVET or TRAPP complexes subunits, highlighting their differential involvement of ORF3a and ORF3b in SARS-CoV-2 pathogenesis. Amongst noticeable proximal interactors, ORF3b identified: RalA, LYN, ALCAM and TMF1. RalA (Ras-related protein Ral-A) is a GTPase involved in multiple mechanisms, including membrane trafficking, exocytosis and filipodium assembly^97^. ALCAM (Activated leukocyte cell adhesion molecule, CD166) is a cell-cell adherence membrane protein contacting CD6 on T-cells, thus contributing in forming the immunological synapse. It has also been reported to mediate dendritic cell attachment to endothelial cells through homotypic interactions^98^. TMF1 (TATA element modulatory factor 1) has been reported to mediate STAT3 degradation and could be involved in RAB6-dependent vesicular transport from endosome to Golgi and from Golgi to ER^99^. Upon poly(I:C) pre-treatment, ORF3b gained several interactors, including USP11 which displays an inhibitory activity on the NFκB pathway^100^. Interestingly, stimulating the innate immune response resulted in a massive loss of ORF3b interaction, suggesting that ORF3b alone is not enough to counteract the antiviral response, which seems to disrupt its function.

**Figure 20.**
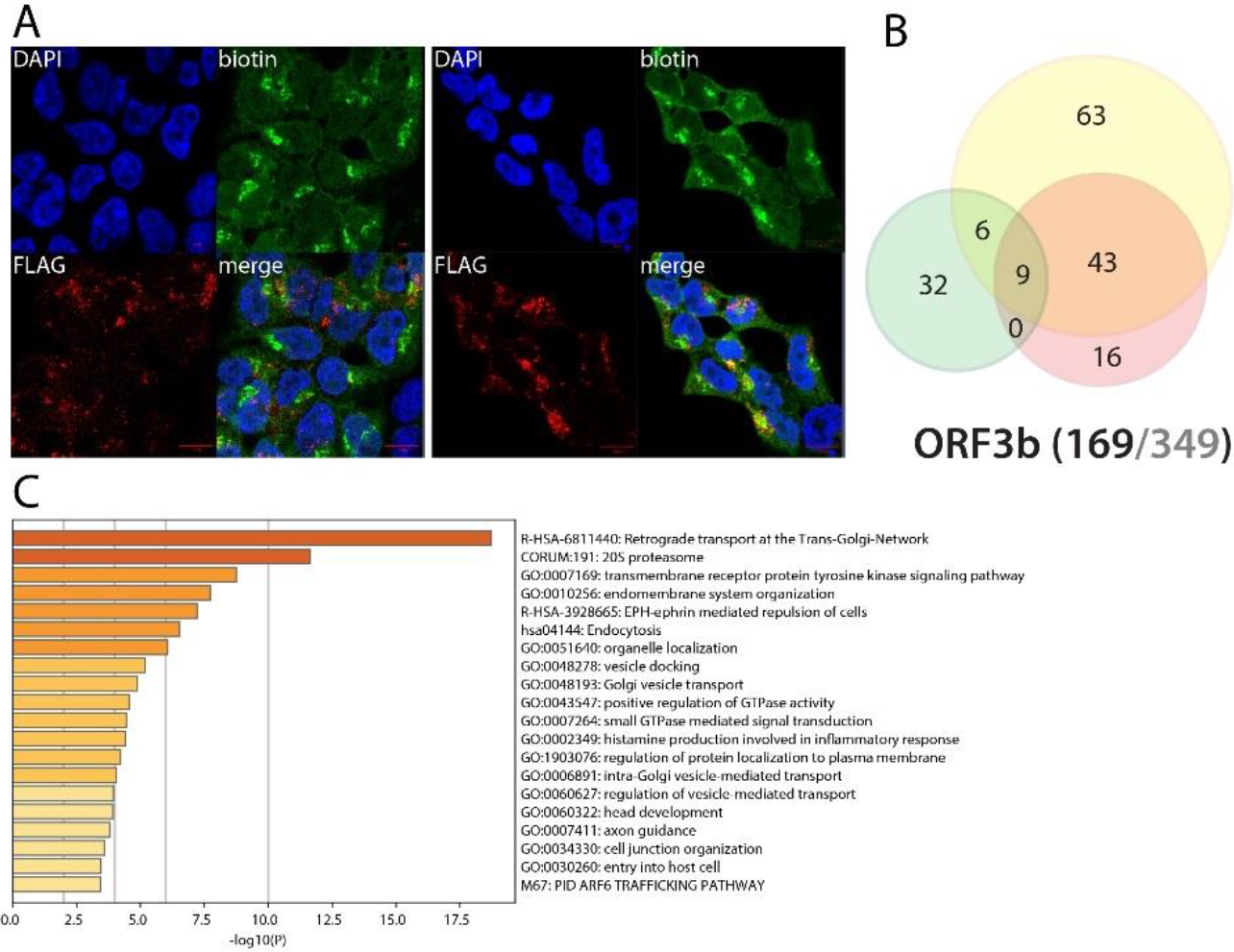
ORF3b summary. See legend Figure 1 (imaging of BirA *Flag-ORF3b).

### Envelope (E) is associated with lipid metabolic pathways and might regulate lipid source for virion assembly

With Spike (S) and Membrane (M) proteins, E is essential to virus particle structure and contributes to the interface between the virus and the environment. Prior to virion formation, structural proteins are expressed into infected cells along the viral cycle. During SARS-CoV-1 cycle, a large amount of the very similar E protein is expressed into host cells, but only a small portion is incorporated into the virion envelope^101^, strongly suggesting non-structural roles of E in hijacking host cell mechanisms. E is a short transmembrane protein localized at the site of intracellular trafficking (ER, Golgi, ER-Golgi intermediate compartment -ERGIC). Expectedly, given its strong membrane association, we did not detect any overlapping interactors with previous reports^1,2,3^. Most E proximal interactors were localized at the ER, ERGIC or Golgi membranes. Other categories such as lipid metabolic process or ion membrane transport were also highly enriched. Interactors associated to *e.g*. vacuole and mitochondria suggest that E could actively participate in recruiting membranes and lipid processing machineries at the virion assembly site to provide suitable lipids for viral particle assembly. Amongst 226 identified proteins, BioID revealed high confidence interactions between E and: C1QBP (coagulation^102^, inflammation^103^, infection and apoptotic processes^104^), TMEM59 (autophagy^105^), TMEM9B (enhances TNF, IL-1B, IL-6 and TLR ligand production; involved in NFκB and MAPK pathways activation^42^), BTN2A2/3A2/3A3 (IL-2 and type II IFN regulation^106^, inhibits type II IFN (IFNγ) release and T cell regulation^107^, respectively), TGFBR1 (TGFβ main receptor, involved in cell homeostasis and apoptosis regulation^108^). E also detected KCNJ8/11, two subunits of the ATP-sensitive potassium channels (KATP) which is involved in cardiac and smooth muscle regulation (see^109^ for review). Together, our data suggest that E plays a role in recruiting fatty acids sources for viral assembly, and could impact ceramide, cholesterol and glycosylation pathways to make them suitable for virion envelope constraints. The putative links between E and coagulation (C1QBP) and cardiac muscle regulation proteins (KCNJ8/11) is also of interest considering the atypical COVID-19 frostbite-like skin rashes. Poly(I:C) pretreatment induced gain of interactions such as ZC3HAV1 (ZAP), involved in viral RNA degradation and a regulation of RIG-I downstream signaling^38,39,110^, suggesting a putative role of E in counteracting host antiviral defense mechanisms.

**Figure 21.**
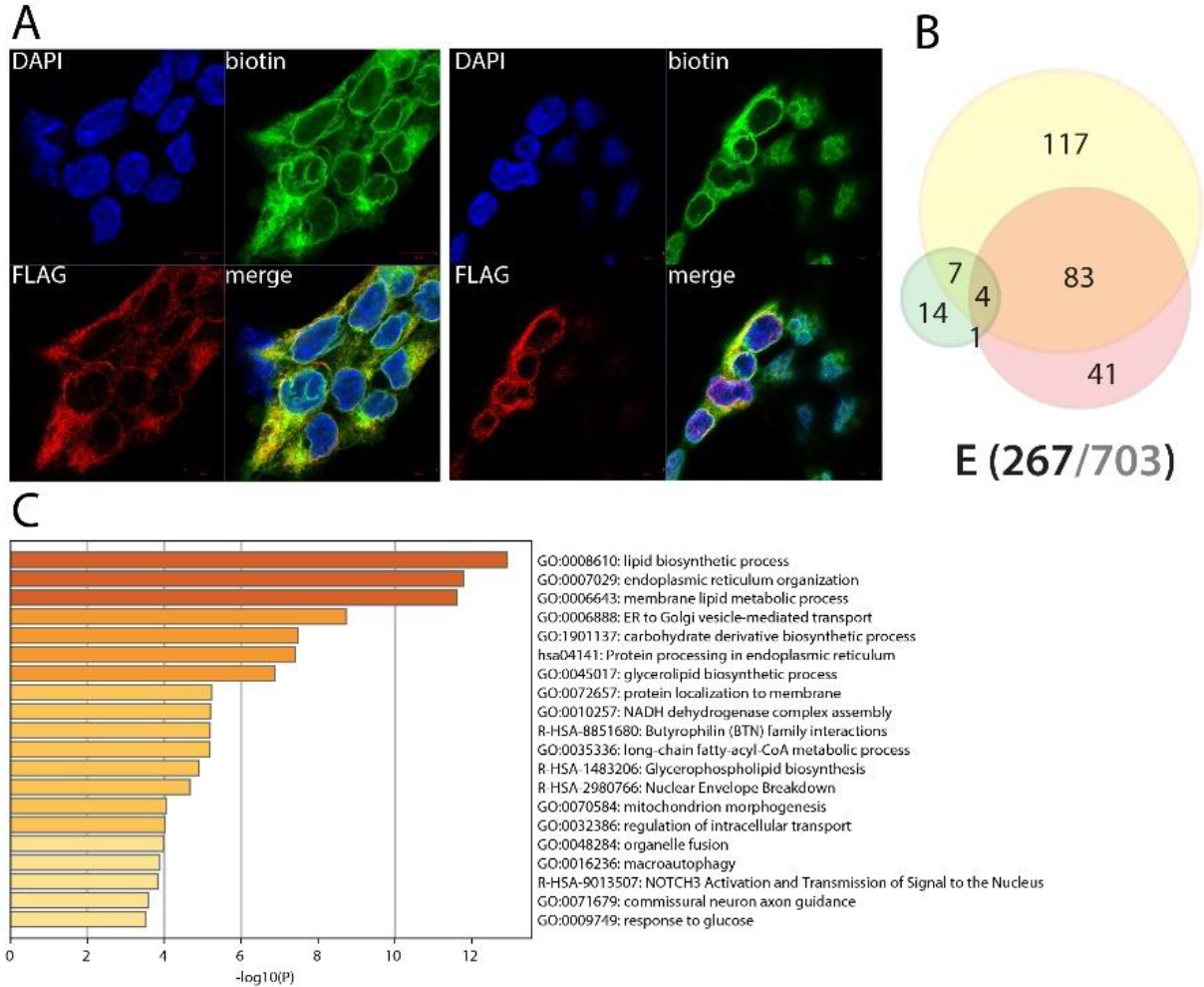
Envelope (E) summary. See legend Figure 1 (imaging of BirA*Flag-E).

### Membrane (M) interacts with membrane shaping factors at the ER or Golgi and capture innate immune pathways regulators

The SARS-CoV-2 membrane protein (M) is a structural component of the viral particle, and through interacting with the other structural proteins S, E and N, plays a major role in virus morphogenesis and assembly. By homology with SARS-CoV-1, M is localized at the ER, ERGIC and Golgi and is thought to be involved in NFκB signaling^111^ and apoptosis^112^. We identified 10 previously reported interactors of Our M BioID data are highly enriched in membrane proteins (ER, Golgi, ERGIC, plasma membrane). Given the complexity of M proximal interactome and to narrow down our analysis to groups of interactors likely to be related to M functions, we decided to focus on M interactions detected by at most three viral bait proteins. Amongst these 198 highly specific interactors, we identified *e.g.:* 37 cell junction proteins; 32 lipid binding proteins; 12 hits related to actin cytoskeleton organization; 14 proteins regulating phosphatidylinositol metabolism; four components of the AP-1 complex (AP1B1/G1/M1/S2); ten components regulating the MAPK activity. These interactions already suggest a much wider involvement of M in the SARS-CoV-2 viral life cycle than suspected so far. We also detected highly specific interactors (detected by ≤3 viral bait proteins) potentially linking M to COVID-19 related processes, such as: PRKD2, ERBB2IP, IL6ST, TBC1D23, B2M, ADD1/2/3, EHD1, EHD4, SNXs, FCHO2, and SH3GL3. The Serine/threonine-protein kinase D2 (PRKD2) is a regulator of the MAPK, NFκB and cytokine production pathways^113^. ERBB2IP (Erbb2-interacting protein) inhibits NOD2-dependent NFκB signaling and proinflammatory cytokine secretion^114^. TBC1D23 (TBC1 domain family member 23, also known as HCV non-structural protein 4A-transactivated protein 1) may act as a general inhibitor of innate immunity signaling, strongly inhibiting multiple TLR and dectin/CLEC7A-signaling pathways^115^. B2M (Beta-2-microglobulin) is a component of the MHC-I and is thus involved in antigen peptide presentation^116^. ADD1/2 are alpha- and beta-adducin, respectively, and are responsible of membrane-cytoskeleton binding and involved in spectrin-actin assembly^117^. EHD1 and EHD4 (EH domain-containing protein 1 and 4) control membrane tubulation in an ATP hydrolysis-dependent manner^118^. SNX1/2 (Sorting nexins 1 and 2) are involved in retromer transport and in sensing membrane curvature^119,120^. FCHO2 (F-BAR domain only protein 2) can recruit membranes and modify their curvature as well^121^. Finally, SH3GL3 (Endophilin A3) has been reported to recruit other proteins to membranes with high curvature^122^. Together, these data show that M can impact diverse pathways, such as MAPK, NFκB, innate immunity, antigen presentation and membrane curvature regulation. This later process is of interest since M is known to regulate virion membrane shaping^123^. Membrane curvature depends of lipid composition and a nexus of regulation involving proteins such as those cited beforehand (containing a BAR-domain to regulate curvature), cytoskeleton-membrane attachment proteins (ADD1/2) to push membranes, form necks and allow budding. Upon poly(I:C) induction, specific interactors were gained including five heat shock proteins, two components of the ESCRT-0 complex, and three BAG-domain containing proteins (BAG2/4/5). The other noticeable interactors were IFIT1 and TAP2 (Peptide transporter involved in antigen processing 2), a protein involved in the assembly of the antigen peptide to the MHC-I. Along with TAP1 (detected in basal culture conditions), it forms the tapasin complex, which is targeted by multiple viruses as a strategy to impair antigen peptide presentation and escape immune system activation^124,125^. Our data suggest that M recruits specific membrane remodeling components of the host machinery to allow virion formation. It could also be involved in the regulation of the pathways mentioned hereabove.

**Figure 22.**
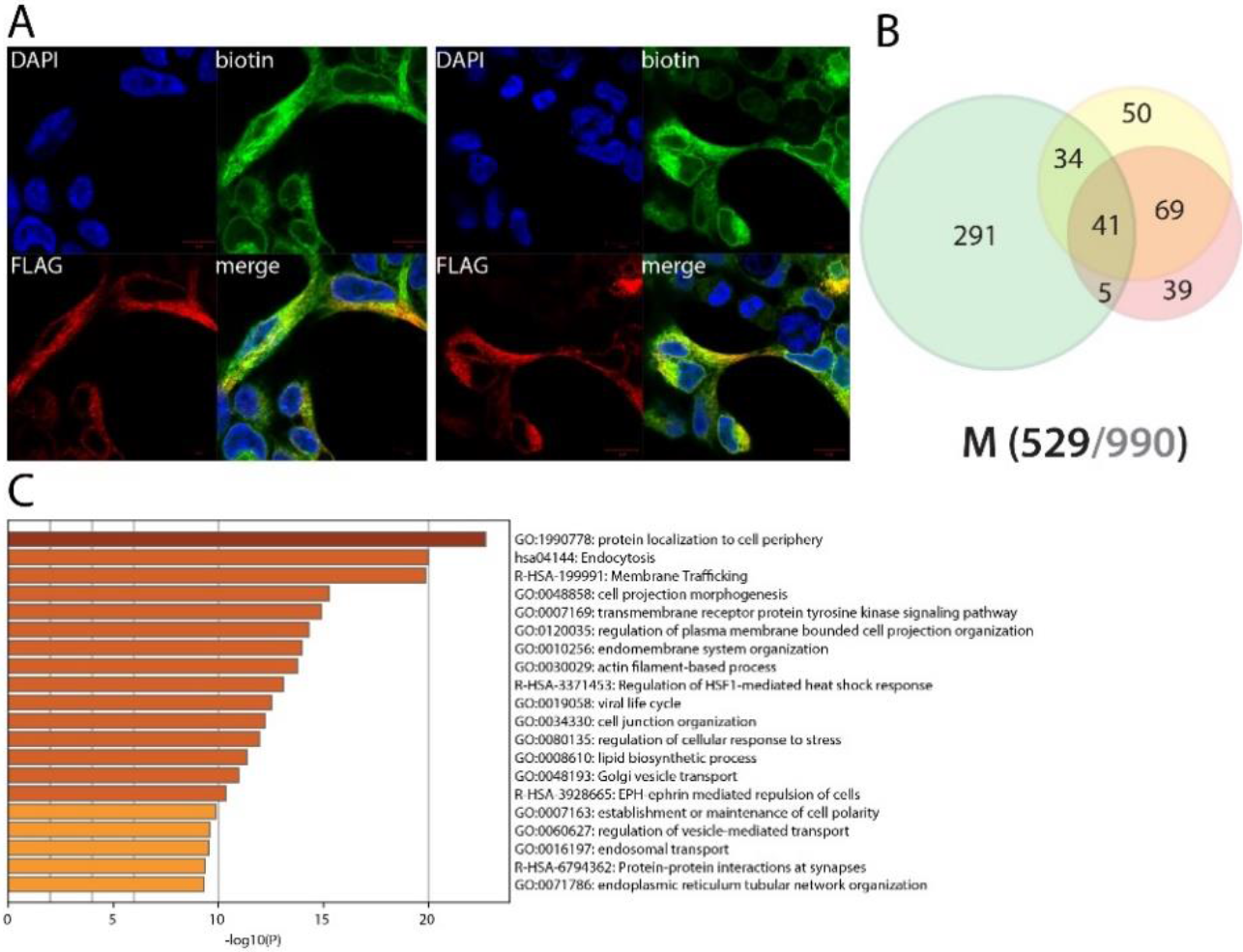
Membrane (M) summary. See legend Figure 1 (imaging of BirA*Flag-M). M^1,2,3^

### ORF6 uniquely captures the CCC complex, several components of the nucleopore inner channel, and could thus impair host cell proper innate immune response

ORF6 is an accessory protein shown to inhibit IFN signaling through disrupting karyopherin-dependent nuclear translocation of transcription factors such as STAT1 (by homology with SARS-CoV-1)^126,127^. A recent study also characterizes SARS-CoV-2 ORF6 as a potent inhibitor of IFNβ and NFκB pathways and IFN stimulated genes such as ISG54 and ISG56, which together, make ORF6 a major antagonist of innate immune response^128^. Unexpectedly, our BioID experiments did not identified ORF6 specific high confidence interactors when the tag was positioned at the C-terminal end. We identified four previously reported interactors of ORF6 (DCTN2, NUP98, RAE1 and KPNA2). KPNA2 has been previously associated with SARS-CoV-1 ORF6 immune escape mechanism^126,127^, impairing STAT1 nuclear import. ORF6 BioID identified different interactor groups: 18 components of the nucleopore, including four out of six subunits of the central transport channel (NUP35/54/62/214); six out of ten COMM-domain containing proteins (COMMD1/2/3/4/6/8), associated with CCDC22 and CCDC93, which together form the CCC complex involved in LDL cholesterol levels and membrane recycling^129,130^; 11 centrosomal proteins including four centriolar satellite factors (PCM1, OFD1, CEP131, SPAG5); and four interactors assigned to the COPII-coated ER to Golgi transport vesicle (SEC13/23A/24C/31A). The interactions linking ORF6 to the nucleopore complex are unique across the 28 viral bait protein interactomes. Indeed, this is the only bait to identify two out of the four central channel factors detected (NUP62/214), which is in line with the CoV-1 ORF6 reported function of impairing transcription factors nuclear import^126,127^. Likewise, ORF6 was the sole bait to detect the CCC complex components. Besides its reported role on cholesterol, several CCC complex components have been shown to downregulate NFκB, promoting its ubiquitination and subsequent degradation^131^. ORF6 identified several peroxisomal (ABCD3, PEX19, PEX11B) and mitochondrial outer membrane proteins (TOMM70A, USP30, ARMCX3, MFF, DNML1, AKAP1, BCL2L13). In addition, ORF6 identified additional factors which could also contribute to its antagonizing role on innate immune response: TBK1 and MAVS, the antiviral signaling adaptor protein downstream RIG-I and MDA5^132,133^. Induction of IFN response through poly(I:C) induced the ORF6-TRIM56 proximal interaction, which could provide another key regulation layer on innate immune response. It also induces interactions with five 26s proteasome components, suggesting a putative involvement of ORF6 in targeting innate immune response factors to degradation. Taken together, our data show that ORF6 interacts with specific components of the nucleopore complex, probably impairing nuclear translocation of transcription factors driving innate immune response program, and uniquely identify the CCC complex, which can target NFκB for degradation. The interaction with TBK1, MAVS and TRIM56 could indicate another level of regulation, given their action on the early stages of innate immune pathways activation (downstream RIG-I, MDA5, TLR3, STING).

**Figure 23.**
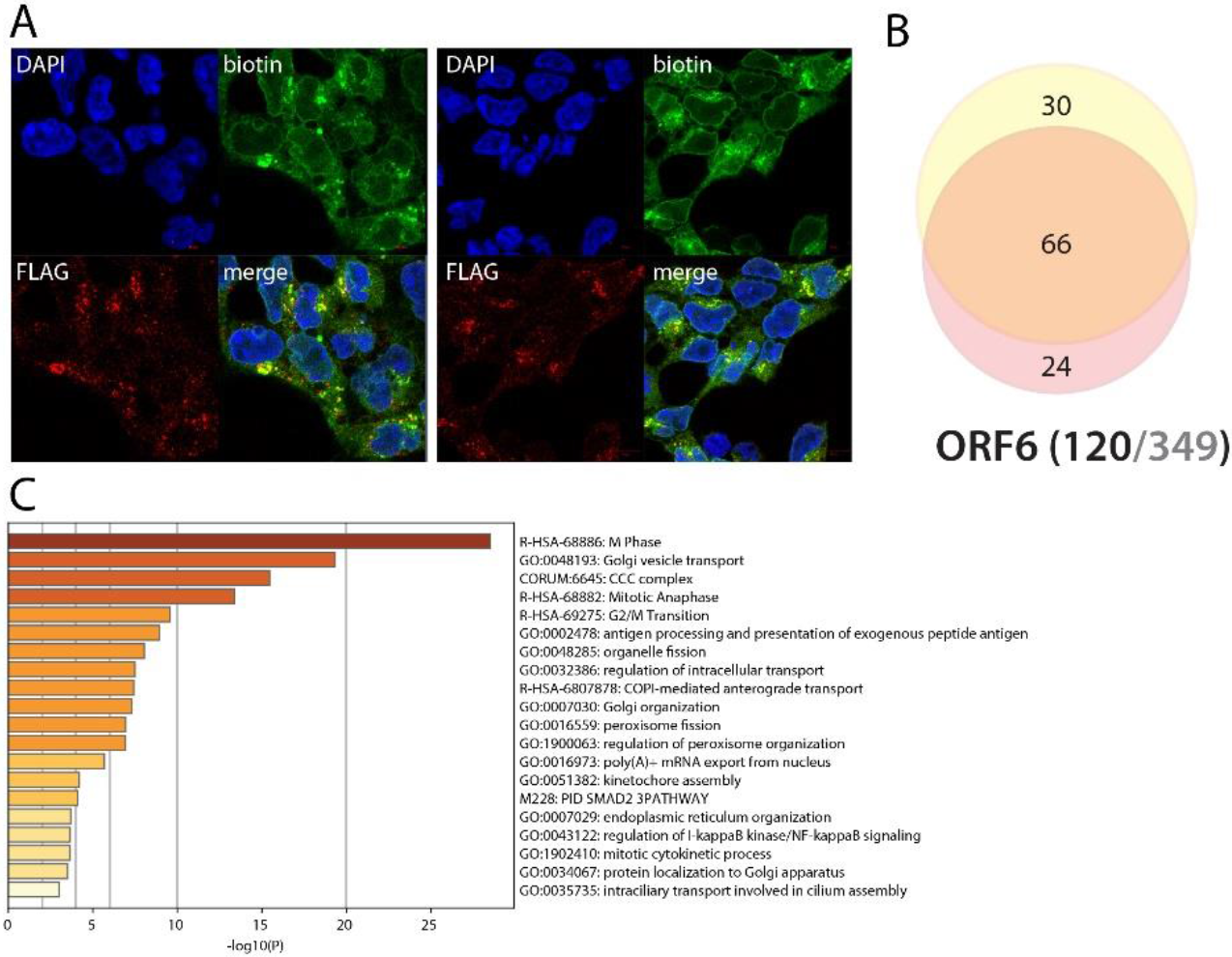
ORF6 summary. See legend Figure 1 (imaging of BirA *Flag-ORF6).

### ORF7a is a membrane-associated protein potentially interfering with innate and adaptative immune pathways, coagulation regulation and olfactory receptors homeostasis

Little is known about the SARS-CoV-2 ORF7a accessory protein. The SARS-CoV-1 ORF7a has been reported to antagonize the host restriction factor BST-2/Tetherin and to act as an apoptosis inducer^134,135^. We identified four reported ORF7a interactors: FANCI, UBE3C, NUP205 and NDUFA12. ORF7a BioID analysis revealed that 282/329 proximal interactors were associated with membranes (129 ER, 97 vesicular, 67 Golgi and 129 plasma membranes). Expectedly, ORF7a interactors are involved in ER, Golgi, vesicles and membranes-related processes, such as lipid biosynthetic processes (42), transmembrane transport (58), protein N-linked glycosylation (13), secretion (35), or antigen processing and presentation (12). Amongst ORF7a proximal interactors linked to innate or adaptative immune responses, we identified *e.g*. UNC93B1^36^(TLR3/7/9 endosome addressing^136^); BTN3A2/3A3 (T-cell response and IFNγ release inhibition); HLA-B, B2M, TAP2 (antigen processing); NCSTN, APH1A, OSTC, BACE2, TMEM59/59L (Notch and APP related pathways); CD97 (leukocyte adhesion and activation^137^); PRKCQ (IL-2 and NFκB pathways); SPPL2B (intramembrane cleaving protease; ITM2B^138^ and TNF processing^139^); IKBIP (NFκB pathway); NPLOC4-VCP-UFD1L complex (type I IFN inhibitor^44^); TMEM9B (enhancer of proinflammatory cytokines in response to IL1β, TNF and TLR activation; activator of NFκB and MAPK pathways through TNF stimulation). We also noticed the presence of 6/12 components of the OST-complex (STT3A, DDOST, OSTC, DAD1, TMEM258, RPN2). GGCX (Vitamin K-dependent gamma-carboxylase) is another host-factor of interest, since it could potentially be linked to coagulation dysfunctions observed in COVID-19^140,141^. In addition to these proximal interactions of interest, we identified several factors involved in processes potentially related to anosmia and ageusia symptoms^142^. ORF7a identified the REEP proteins which are involved in membrane receptors and/or olfactory receptors regulation mechanisms^143^. Along with ubiquitin-proteasome system interactions, an attractive hypothesis would be that ORF7a drives an olfactory receptor membrane addressing/recycling impairment through proteolysis or sequestration of one or several REEP protein(s). Together, our data reveal an ORF7a complex nexus of proximal partners potentially involving this accessory protein in a broad range of mechanisms, including yet poorly understood pathophysiological mechanisms (hyper clotting, anosmia, ageusia) related to COVID-19. Upon poly(I:C) induction, ORF7a gained several interactors including TRIM4, an E3 ubiquitin ligase targeting RIG-I for K63 polyubiquitination, which leads to an augmented IFN signaling activity^144^.

**Figure 24.**
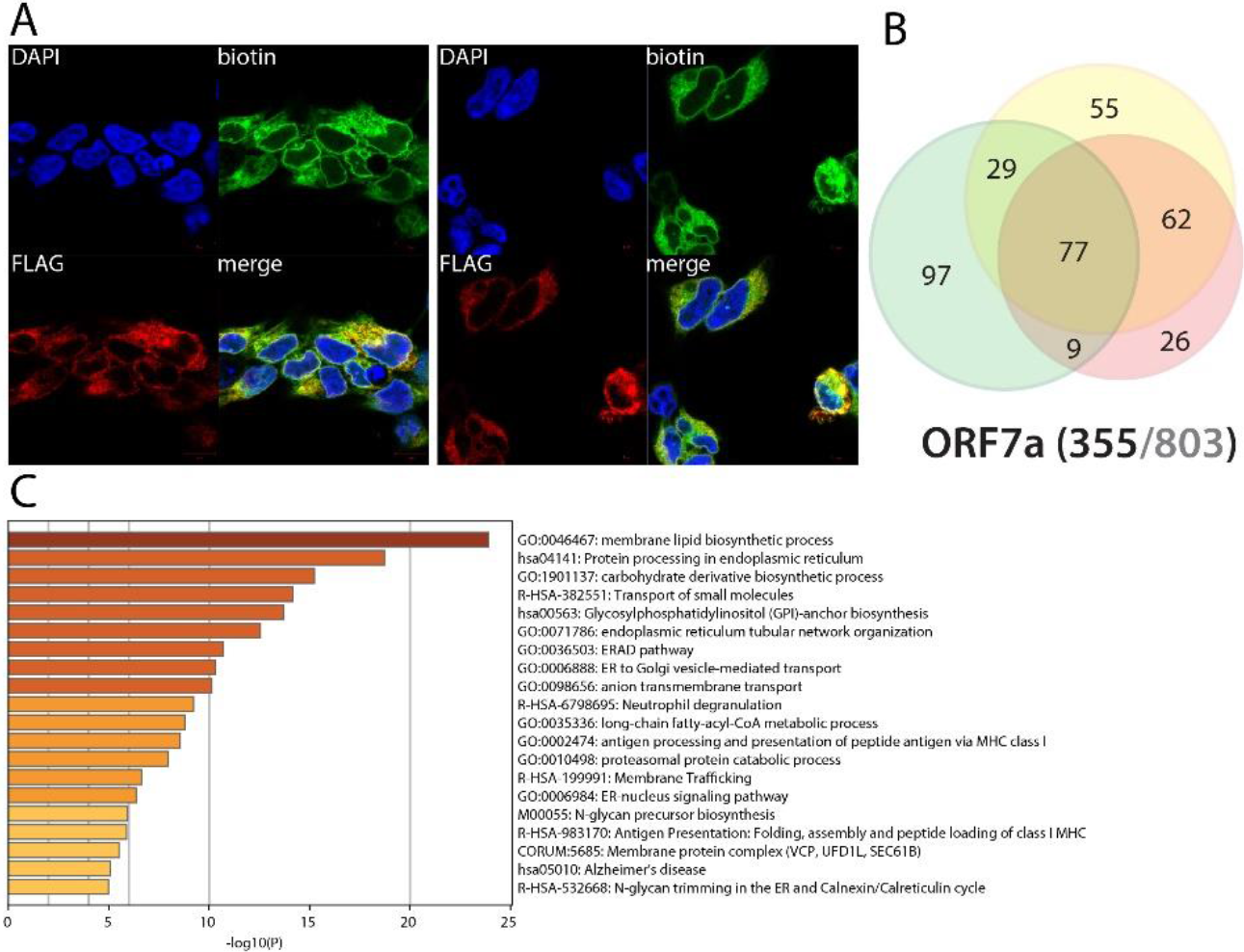
ORF7α summary. See legend Figure 1 (imaging of BirA *Flag-ORF7α).

### ORF7b is likely to cooperate with ORF7a in a multiple membrane-based processes, ranging from cell adhesion, olfaction and viral life cycle

To date, there is no functional information available on SARS-CoV-2 ORF7b. It is retained at the Golgi apparatus and has been shown to be 5) incorporated into the viral particle in SARS-CoV-1^145^. Comparing our data with previously reported ORF7b interactions, we identified 85 proteins in common, including vesicle docking/membrane fusion factors (*e.g*. STXs) and important inflammatory and antiviral pathways regulators such as ITCH, UNC93B1 and IL6ST. About half of ORF7b proximal interactors were shared with ORF7a, suggesting these two viral proteins could cooperate in several mechanisms (*e.g*. lipid metabolism, immune pathways regulation, vesicular recycling of membrane receptors, antigen presentation). ORF7b differentially interacts with 232 hits (compared to ORF7a) highly enriched in cell adhesion (32). Regarding viral infection-related proximal interactors, ORF7b identified: IL6ST; LIFR (signal transduction; type I cytokine receptor, potentially sharing pathways with IL6ST^146^); IL1RAP (interleukin-1 receptor accessory protein); STAM, STAM2, HGS (ESCRT-0 complex; IL-2 and GM-CSF cytokines signal transmission in association with STAM); ALCAM (CD166); LYN; ITCH (inflammatory signaling pathways). Of note, we detected furin uniquely interacting with ORF7b. Furin plays a role in S protein-driven membrane fusion process upon early stages of SARS-CoV-2 entry^147^. Regarding anosmia symptom, we detected GRID1, the glutamate receptor ionotropic, delta-1, which is essential for olfactory cells signal transmission^148^. In summary, ORF7b seems involved in similar mechanisms as ORF7a, but targets different factors. It seems more specialized in cell adhesion and could act on different components of olfactory signaling cascades. Following poly(I:C) treatment, ORF7b interactome is greatly impacted with 91 interactors increased/gained (including *e.g*. GRID1) and 67 decreased/lost, suggesting a regulation upon IFN pathway activation. More than half of these gained hits are assigned to the plasma membrane. Amongst enriched/gained interactors, we identified *e.g*.: ROBO2, which is involved in olfactory bulb neuron development^149^; IFIT1, TOLLIP, BTN3A2/3A3 (immunity); and CD83 (MHC-II antigen presentation^150^). ORF7b thus could be involved in multiple host pathway deregulations, such as inflammation, IFN pathway activation and olfaction.

**Figure 25.**
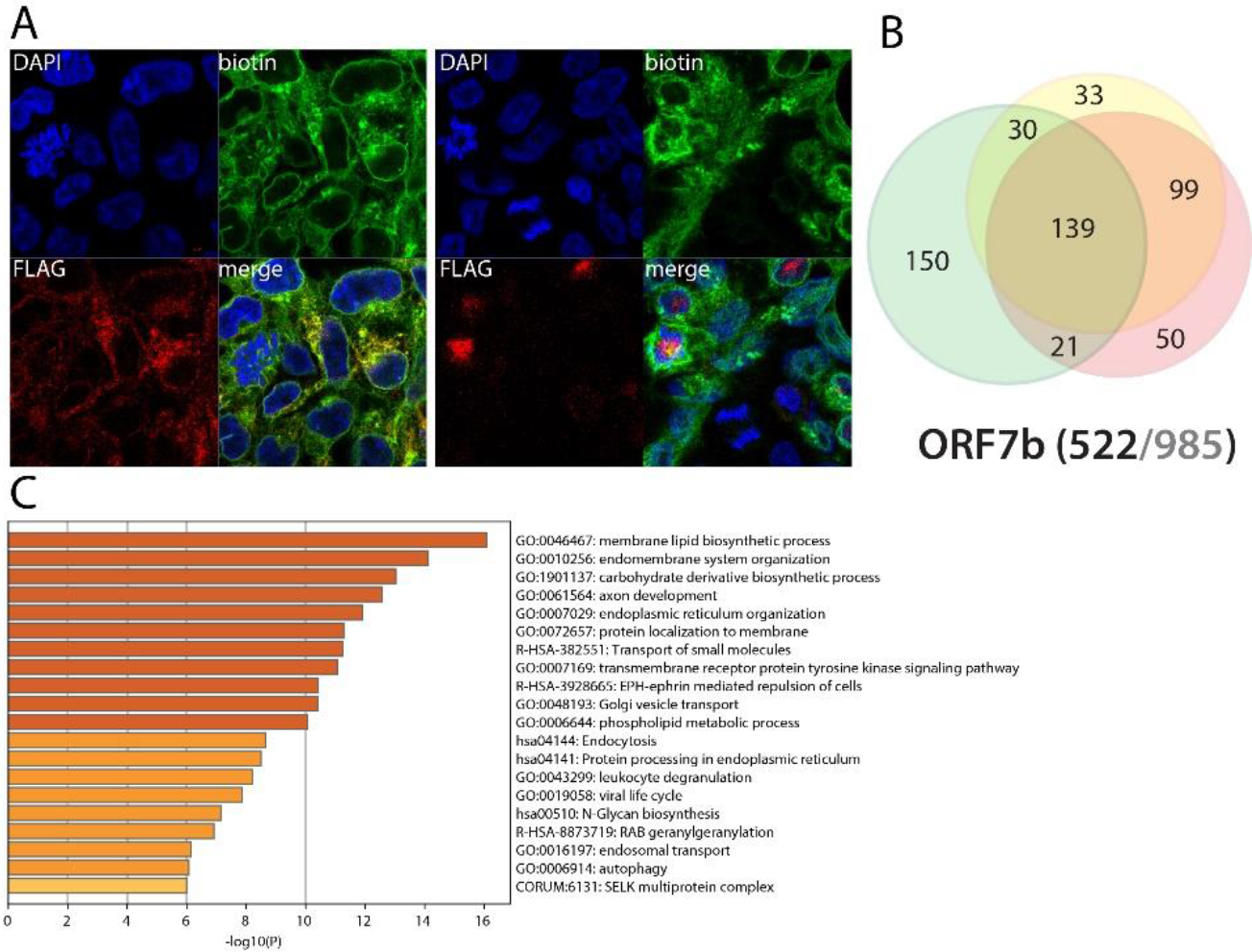
ORF7b summary. See legend Figure 1 (imaging of BirA *Flag-ORF7b).

### ORF8 could be involved in MHC-I peptide presentation, translation and phagocytosis regulation

ORF8 is a SARS-CoV-2 accessory protein showing the lowest homology with SARS-CoV-1-encoded sequences and is supposedly involved in innate immune response through downregulating MHC-I^151^. Looking at highly connected ORF8 interactions, we detected several components potentially linked to MHC-I peptide presentation: HLA-C; BCAP31; SEC22B/24B; SAR1B; TAP1. We also identified eight subunits of the endoplasmic reticulum membrane complex (EMC); six components of the proton-transporting vacuolar (V)-ATPase protein pump (CCDC115, TMEM199, ATP6V0A1/0A2/1F, ATP6AP1); three components involved in platelet calcium metabolism (ITPR1/2/3). ORF8 interactions connected to six or less other proteins revealed 120 specific interactors. Besides 20 hits related to signal recognition particle-dependent co-translational protein targeting to membrane and 21 ribosomal proteins from both small and large ribosomal subunits, we did not detect enriched interactor group worth highlighting. Amongst individual annotation, we can stress a few ORF8 proximal interactors: POGLUT1 (Notch O-glycosylation^152^); MID1 (E3 ligase directing lytic granules exocytosis in cytotoxic lymphocytes^153^); and TRIM2 (ubiquitin ligase restricting New World Arenavirus entry, probably regulating phagocytosis^154^). Upon poly(I:C), among others we gained interaction with IFIT1. Together, the ORF8 BioID interactome did not provide striking clues to narrow down its putative functions in cells.

**Figure 26.**
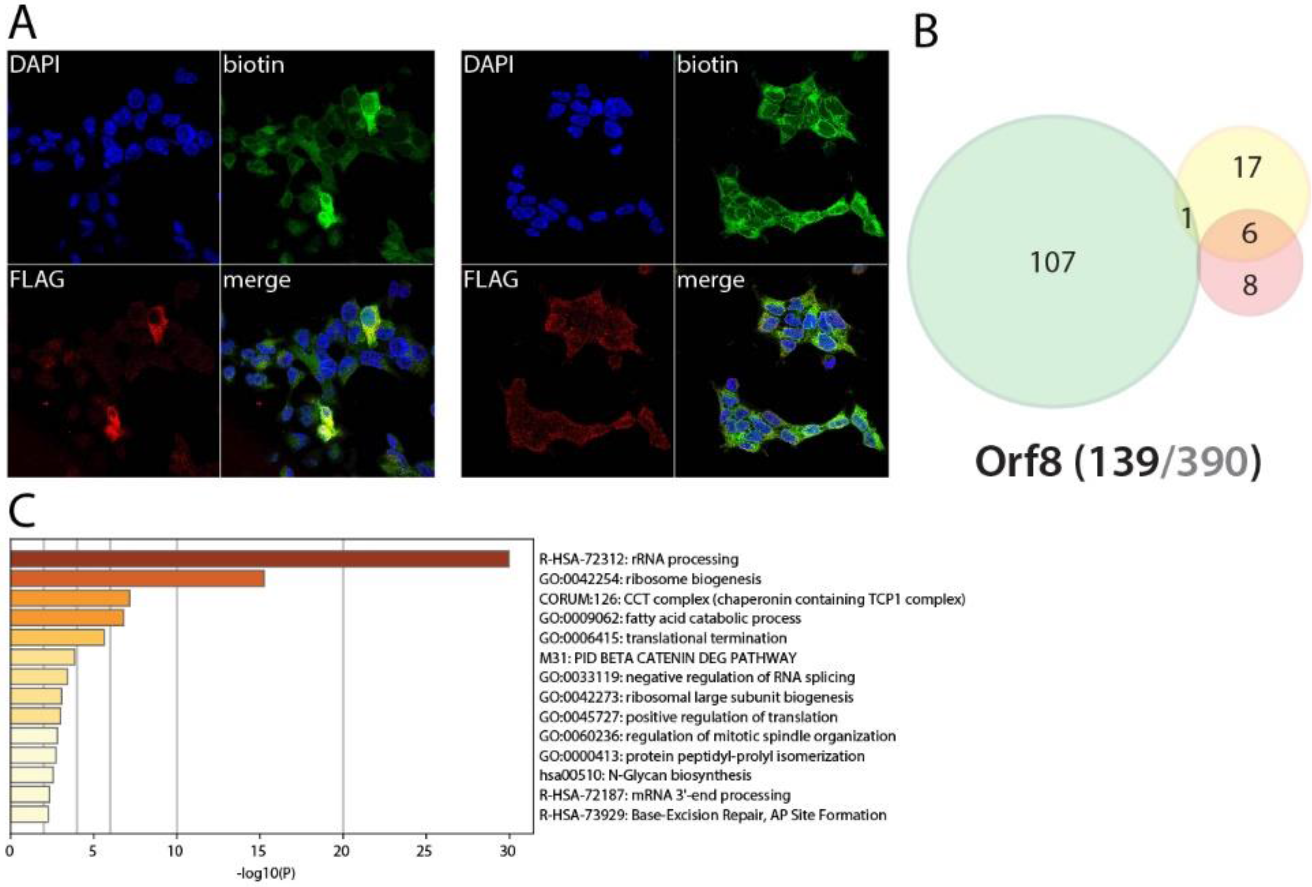
ORF8 summary. See legend Figure 1 (imaging of BirA *Flag-ORF8).

### N interacts with RNA granules components, and upon poly(I:C) stimulation, with PKR, IFIT1/2/3/5, TRIM26 and appears as a key regulator of host innate immune response

SARS-CoV-2 Nucleoprotein (N) is a structural protein associated with the viral genome. It is highly immunogenic and abundantly expressed during infection^155,156^. N binds viral RNA through both its N- and C-terminal domains, connected by a disordered SR-rich linkage region^157,158^. Of note, given its high abundance within the viral particle, N is the first viral protein released in the host, prior to viral RNA translation. It is thus expected to achieve “emergency” pro-viral functions to create a cellular environment suitable for viral replication. In line with previous interactomic analysis, we identified 10 reported interactors, including G3BP2, CAVIN1 and GSK3A/B and PRKRA as N high confidence proximal interactors. These data suggest that: N can localize within the stress granules (G3BP2), cytosolic non-membrane organelles involved in mRNA storage; interact with caveolae formation (CAVIN1); is involved in the Wnt pathway and/or glycogen metabolism (GSK3A/B); and could interfere with IFN response (PRKRA, or PACT). Our BioID data also identified eleven RNA or stress granules associated proteins: XRN1 (nucleases); ZCCHC6 (or TUT7, miRNA regulation); TNRC6B and DICER1 (RNA-mediated gene silencing machinery cofactors); STAU2; G3BP1; TDRD3 (RNA granules); FAM120A, CPEB4, PTRF, SRP68, SECISBP2 (RNA transport and/or binding). This is in line with the reported property of N to phase separate with RNA^159^.Importantly, N interacts with factors involved in antiviral host response, such as TRIM26, an E3 ubiquitin ligase targeting IRF3 for proteasomal degradation and bridging TBK1 and NEMO during infection^160^. In basal condition, N is thus able to interact with RNA binding and processing host-factors, including several stress granules biogenesis factors. We see these interactions as a viral strategy to either protect vRNA from host nucleases, or to control vRNA detection by pattern recognition receptors (PRRs), or to regulate vRNA transcription/replication/translation. The interactions with TRIM26 and PRKRA could reveal N-mediated inhibition of both IFN and NFκB innate immune response pathways. In the poly(I:C)-stimulated condition, N captured additional interactors of outstanding interest: EIF2AK2 (PKR) and its interactors STAU1/2; ANKRD17; TARBP2 and ZNF346; TRIM56; TRIM26; and IFIT1/2/3/5. It appears logical that N, being the first viral protein released in host cells, directly targets these fundamental innate immune response proteins. In SARS-CoV-1, N was found to inhibit IFN production interfering with TRIM25, an Ub E3 ligase mediating RIG-I degradation^161^. Our data suggest that N SARS-CoV-2 could use additional or different mechanisms, such as interfering with RIG-I stabilization by binding ANKRD17^162^, impairing viral RNA recognition through IFIT1/2/5 inhibition^163^, and/or precluding RIG-I downstream response targeting MAVS or IRF3 (IFIT3, TRIM26), and/or, upstream MAVS, blocking the TRIM56-mediated activation of STING and FAM173A. PKR is an IFN-induced dsRNA-dependent serine/threonine-protein kinase that detects viral dsRNA to promote multiple proinflammatory cytokines production^165^. PKR is also able to block translation^164^. Together, our data strongly support a central role of N in impairing host antiviral response.

**Figure 27.**
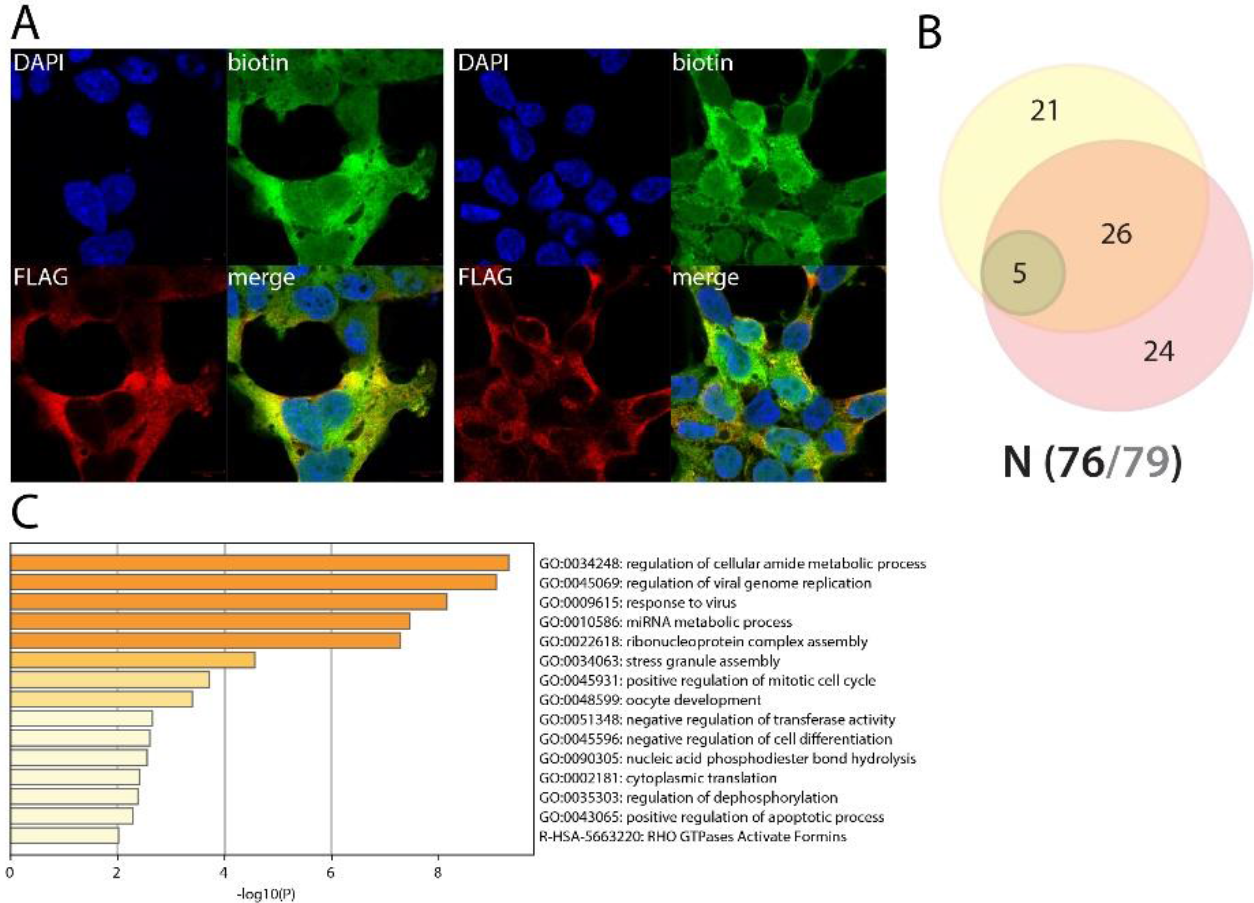
N summary. See legend Figure 1 (imaging of BirA *Flag-N).

### ORF9b interacts with major innate immune pathway factors (MAVS, ISG20) and seems involved in mitochondrial fusion/fission and mitophagy

SARS-CoV-1 ORF9b is encoded by an alternative open reading frame within N and has been linked to innate immune response impairment through its ability to target PCBP2 and AIP4 which leads to MAVS, TRAF3 and TRAF6 degradation^166^. The Kelhr laboratory also reports SARS-CoV-1 ORF9b property of triggering autophagy in an ATG5-dependent manner when highly expressed^166^. ORF9b is located at the outer mitochondrial membrane probably *via* its interaction with TOMM70A^167^. This recent study suggests that the ORF9b-TOMM70A interaction is essential for the type I IFN signaling inhibition role of ORF9b. In our BioID analysis, we identified TOMM70A amongst the top interactors of ORF9b, and MARK1/2/3 and DNM1L which were also previously reported^1,2,3^. Our data greatly expand our current knowledge, identifying 36 mitochondrial components over 63 high confidence interactors. Within this population, we detected remarkable interactors such as MAVS, PPM1A, PPM1B (innate immune signaling), PRKAR2A/2B, USP30, BCL2L13 (mitophagy), MTFR1/1L, MFF, FAM73A and PLD6 (mitochondrial fusion/fission). Our analysis in basal condition is in line with previous results and identifies novel specific interactors likely to be involved in ORF9b-induced mitophagy. Of note, we identified several core regulators of mitochondria fusion/fission, which could be linked to the hyperfused phenotype reported in^166^ when ORF9b is expressed. Together, our BioID analysis provides important insight on the molecular environment of ORF9b at the mitochondrial outer membrane and uncovers specific factors likely to regulate ORF9b-reported processes. Following poly(I:C) pretreatment, we observed a handful of additional interactions with *e.g*. BAX, IFIT2, and ISG20 (Interferon Stimulated Gene 20kDa protein). This latter factor was uniquely detected with ORF9b, and only under poly(I:C) stimulation. ISG20 is an IFN-induced 3’-5’ exonuclease and has an important antiviral impact against multiple viruses, inhibiting their replication probably independently of the exonuclease activity^168^. ORF9b thus appears as a central interference protein of the innate immune response, both targeting the antiviral signal transduction through MAVS, and downstream, the IFN stimulated protein ISG20 to counteract host cell reaction.

**Figure 28.**
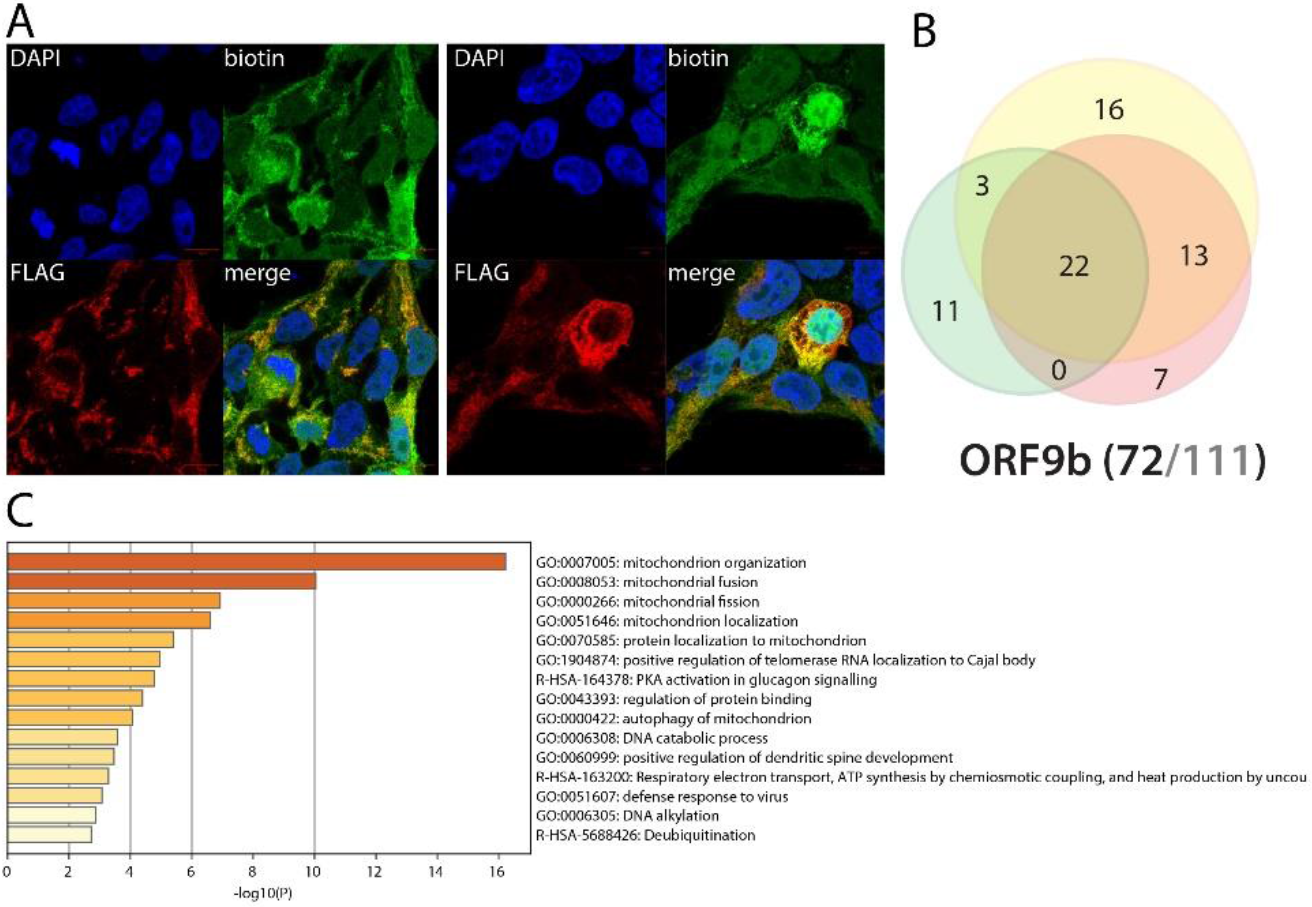
ORF9b summary. See legend Figure 1 (imaging of BirA *Flag-ORF9b).

### ORF14 interacts with rRNA processing pathway components, and chromatin remodeling factors following poly(I:C) treatment

ORF14 (also named ORF9c) is an accessory protein of unknown function encoded by an alternative open reading frame of N, downstream ORF9b^169^. This 73 aa protein has never been studied in SARS-CoV-2, and none of the four reported interactors in SARS-CoV-1 have been identified in our BioID experiments. Whereas the C-ter tagged ORF14 interactome identified proximal interactors mostly localized at the ER and the mitochondrial membrane, the N-ter tagged bait labeled a majority of nucleolar components, which is in accordance with BirA*Flag-ORF14 exclusive localization in the nucleus (**Fig. 29**). This discrepancy indicates either a tag position-dependent targeting signal sterically hindered, or a dual role of ORF14 in these different compartments. The N-ter tagged ORF14 dataset identified 15 components of the preribosome, potentially linking it to the regulation of rRNA processing. Why ORF14 would target the rRNA pathway remains an opened question. Interestingly, several viruses have been shown to impair this pathway to counteract innate immune response^170^. Surprisingly, while the defect of ribosomes produced *de novo* efficiently inhibits the antiviral response, it did not seem to affect viral life cycle. Further investigation must be implemented to test the ribosome production rate in SARS-CoV-2 infected cells. In addition to these interactions, N-ter tagged ORF14 detected USP11 which is involved in NFκB pathway regulation^171^. The C-ter tagged ORF14 interactome includes 37 components of the ER membrane, 35 mitochondrion proteins, four out of five components of the GPI-anchor transamidase complex, and 53 nuclear proteins (including 22 nucleolar). It also identified 11 components of the mitochondrial respiratory chain I, suggesting a role of ORF14 in oxidative phosphorylation. C-ter tagged ORF14 also identified several components of the phagocytic vesicle, including ANXA11, PDIA3, TAP2, RAB8B and UNC93B1. Together, these data suggest that ORF14 uniquely interferes with rRNA processing. In the presence of poly(I:C), N-ter tagged ORF14 gained interactions with 35 nuclear components involved in chromatin organization (16), transcription regulation (23) and DNA methylation (ATRX, EHMT1, BEND3 and BAZ2A). Interestingly, SUMO2 was the most enriched interactor following poly(I:C) treatment. The SUMO system is a major regulator of chromatin compaction^172^. All together, these data suggest that ORF14 could impair chromatin remodeling in response to antiviral pathways activation, hence precluding the induction of innate immune response genes transcription.ORF10 could associate with chaperonin and prefoldin complexes ORF10 is a predicted accessory protein of unknown function^169^. None of the nine previously identified SARS-CoV-1 ORF10 interactors were found in our analysis. BioID identified six chaperonin T complex and five prefoldin complex components, 15 mitochondrial proteins and several ribosomal factors. The presence of poly(I:C) did not induce interaction gain. Overall, the ORF10 interactome did not allow to infer a putative function to this predicted protein. The presence of ORF10 upon SARS-CoV-2 infection remains questionable.

**Figure 29.**
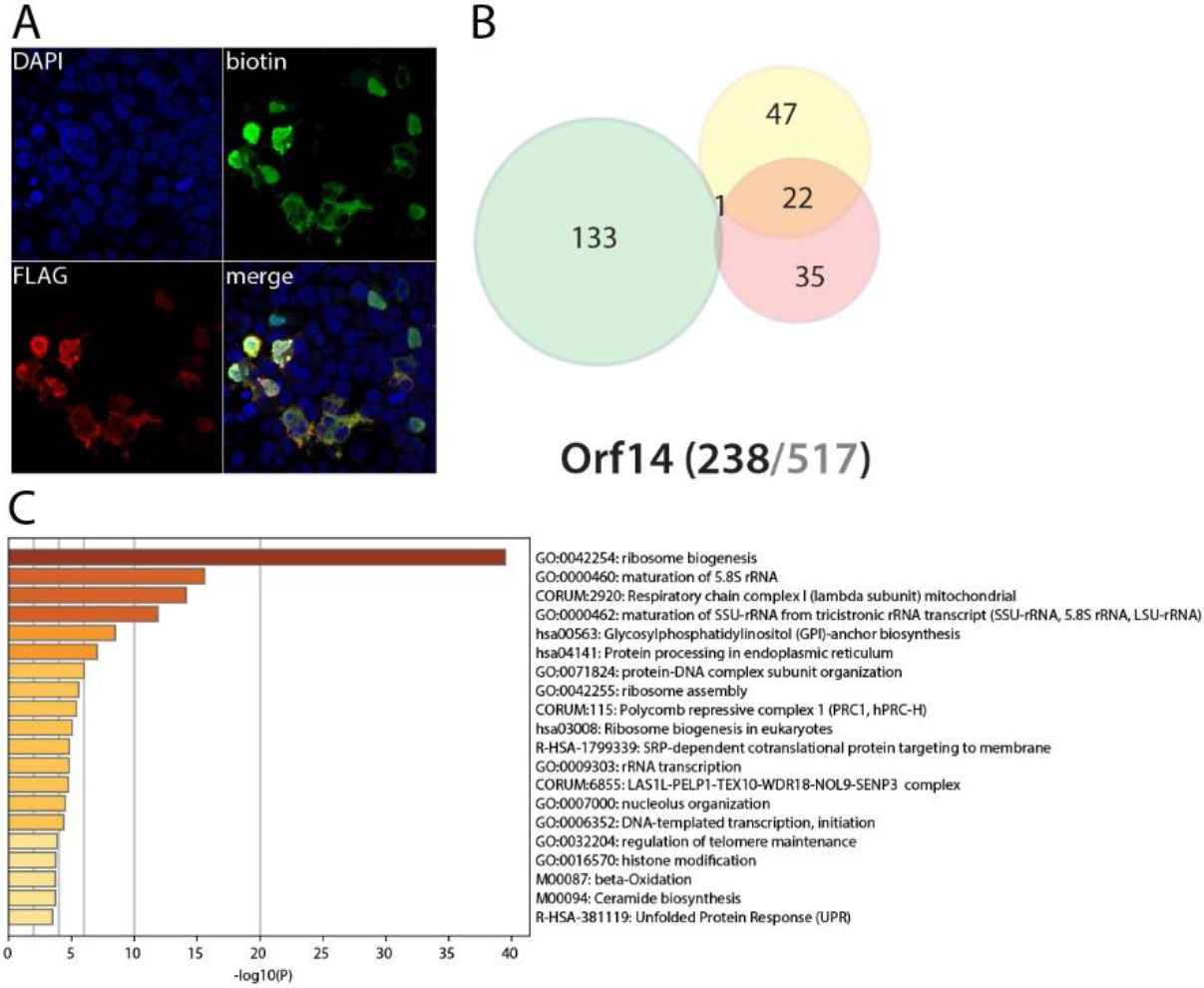
ORF14 summary. See legend Figure 1 (imaging of BirA *Flag-ORF14).

**Figure 30.**
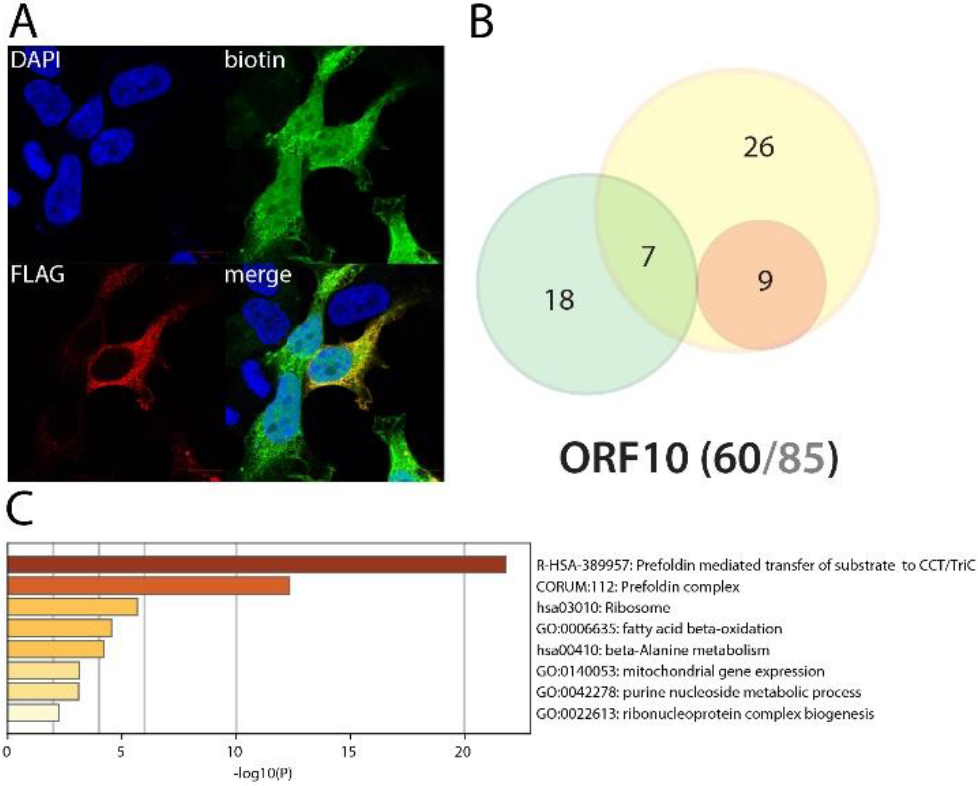
Orf10 summary. See legend Figure 1 (imaging of BirA *Flag-Orf10).

## Discussion

As a general note, the high confidence proximal interactions presented here should be taken with three main cautions: (**i**) our analysis was performed in HEK293 cells, which do not represent a SARS-CoV-2 physiopathological model. However, they are well known to express a broad panel of proteins and are thus excellent bioreactors to identify possible interactions in living cells. In addition, the range of different cell types detected as infected by SARS-CoV-2 keep expanding (*e.g*.^173,174,175,176^). We thus do think that this apparent limitation could be also seen as an asset of our study; (**ii**) In our opinion, the main limitation sits in the expression of a single protein at a time. Knowing that several viral proteins require viral cofactors, infected context and/or the presence viral RNA to function properly (*e.g*. NSP10-NSP14 or NSP10-NSP16), the present analysis almost certainly misses cooperative viral interactions. Similar studies performed in infected cells will thus bring highly valuable additional information on putative SARS-CoV-2 pathogenesis mechanisms. As an attempt to mimic a physiopathological context, we artificially induced an anti-viral response by transfecting poly(I:C) and repeating the proximal interactome analysis. These experiments already revealed novel interactions of the utmost importance; and (**iii**), the proximal interactions are not necessarily physical and should therefore be considered as a discovery step systematically requiring orthogonal or functional validation. However, the proximal interactomics multiple analysis generated by us and others have been at the basis of fundamental mechanism discoveries, supporting the validity of the approach for identifying new biology (see^177^ for review).

This first proximal interaction mapping of SARS-CoV-2 proteins provides a plethora of novel research tracks to better understand this virus pathogenesis. Although for a few proteins our approach did not lead to satisfying results (NSP3, NSP5, NSP8, ORF8 and ORF10), our data clearly assign most of the viral proteins to specific subcellular locations and pathways. Based on our poly(I:C) specific interactomes, we identified NSP2, ORF6, N and ORF9b as the main candidates to explain the defects of innate immune response observed in COVID-19 patients. This study dramatically expands the current knowledge^178^ or uncover novel ties between: NSP1 and translation inhibition mechanisms through eIF3 complex interactions; NSP2 and RIG-I and PARP9-DTX3L, which could inhibit vRNA sensing; NSP4 and inflammation and membrane receptor recycling; NSP6 and lipid metabolism and autophagy; NSP7 and vesicular trafficking; NSP9 and mRNA processing; NSP13 and centrosome; NSP14 and RNA decapping and deadenylation; NSP15 and TLR signaling, mRNA decay and apoptosis; NSP16 and transcription factors nuclear translocation; Spike and cell adhesion and migration; ORF3a and ORF3b and vesicular sorting and fusion; E and lipid metabolism and membrane trafficking; M with membrane curvature and innate immune response regulation; ORF7a and ORF7b and membrane protein recycling, potentially including olfactory receptors recycling, innate and adaptative immune pathways, and vitamin K regulator; ORF7b in cell adhesion and viral life cycle; ORF8 and phagocytic vesicle; N and stress granules and major components of the innate immune response; and ORF9b with antiviral signaling upstream and downstream the IFN-I pathway; ORF14 with rRNA metabolism.

This ongoing work will be subjected to several follow up studies to confirm the role of selected SARS-CoV-2 proteins in specific infectious mechanisms. Our interactome will be subjected to cross validation by orthogonal approaches and mined to identify putative drug targets candidates.

## Acknowledgements

We would like to thank Antoine Touzé, Yves Gruel and Mathias Faure for sharing their invaluable expertise in different SARS-CoV-2 related processes. Soulaimane Aboulouard kindly provided us analysis advices and method materials. We would also warmly thank Irène Gadotti for her outstanding administrative support during the lockdown as well as Nathalie Vasseur for her logistic assistance. Dushyandi Rajendran provided technical assistance to obtain the BioID vectors. Joseph Dias from the Enterprise car rental company provided us a car free of charge to support our research effort facilitating our commute to the lab. Several companies’ sale managers were extremely helpful to facilitate reagents supplies during the lockdown: Karim Haded and Priscilla Vergati (Sigma), Claire Allas-Boone (Sarstedt), Karine Lemarchand (TebuBio), Anne-Lyse Parisot (NEB) and Xavier Flandres (Fisher Scientific). Finally, we would like to warmly thank our supportive co-workers who helped us with their positive attitude and warm encouragements.

## Authors’ contributions

E.MN.L, P.S., DK.K, H.A., J.J.K, D.K. A.C., D.S., A.R., R.L. O.P., AC.G., E.C., Y.J. and F.K.R. obtained and cloned the SARS-CoV-2 polypeptides coding sequences. E. MN. L. produced BioID vectors and conducted all BioID sample purification. E.C. realized the cell culture-related experiments. JP.G. ensured all mass spectrometry acquisition and contributed to the data analyses conducted by E.C.. M.D. assisted in acquiring the microscopy images. D.E.H., M.A.C., P.F-B., P.A., U.S., M.V., AC.G., S.VD.W., Y.J., E.C. and F.P.R. and contributed to the project development through the *VirHostOme* consortium assembled *ad hoc* by F.P.R. to aggregate international interactomics research efforts. I.F. and M.S. supported the project through logistic and scientific contributions. E.C., Y.J., C.D., Y.S., E. MN. L and A.K conceived the project. C.D. and E.C. secured funding. Y.S. conceived the online 3D network interactive visualization interface, with inputs from E.C. and Y.J. and support of G.A.P.. E.C. assembled the figures and wrote the paper, with scientific and editorial inputs from A.K., Y.J., P.C., P.F-B. and E. MN. L.

---

**Figure S1.**
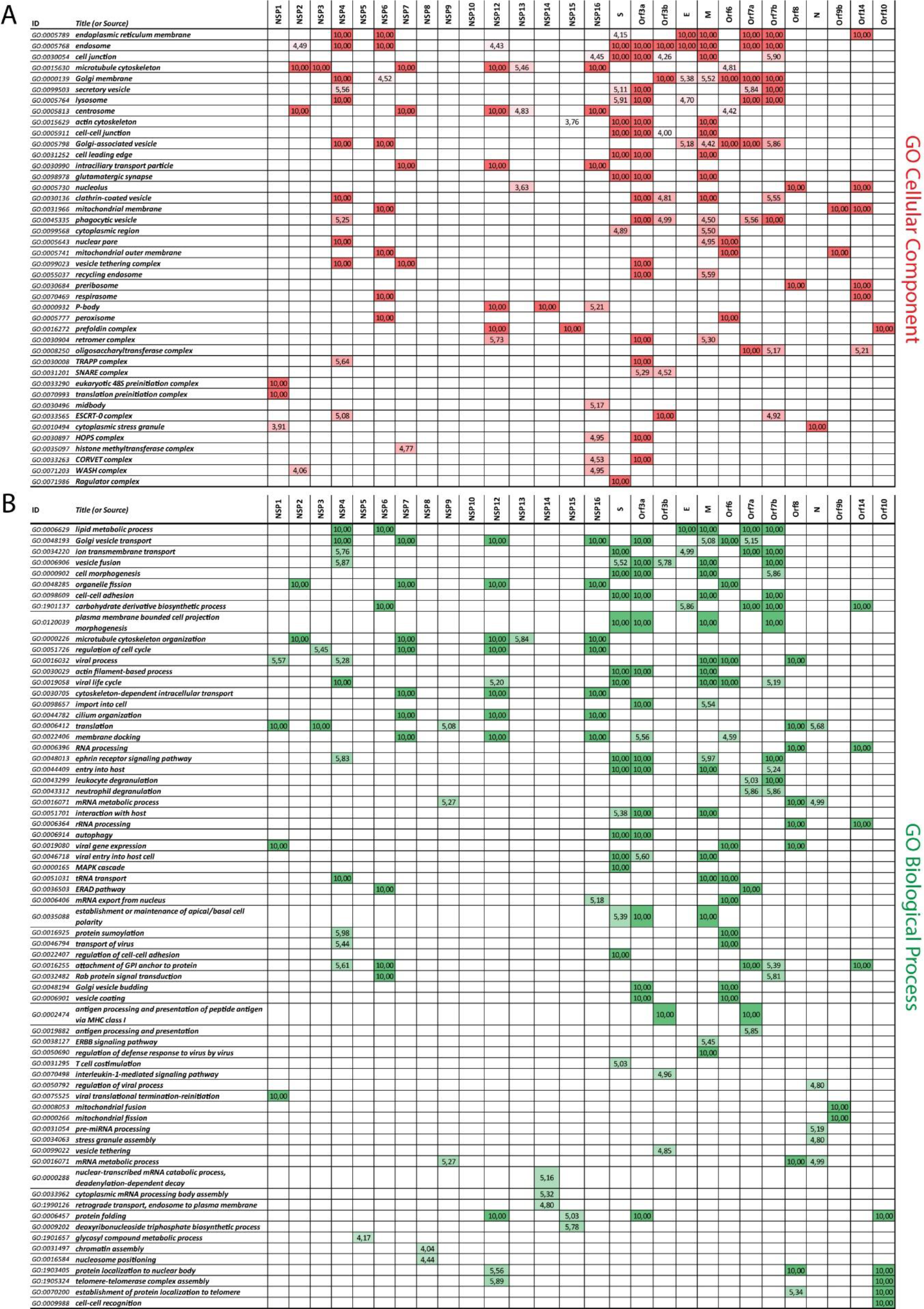
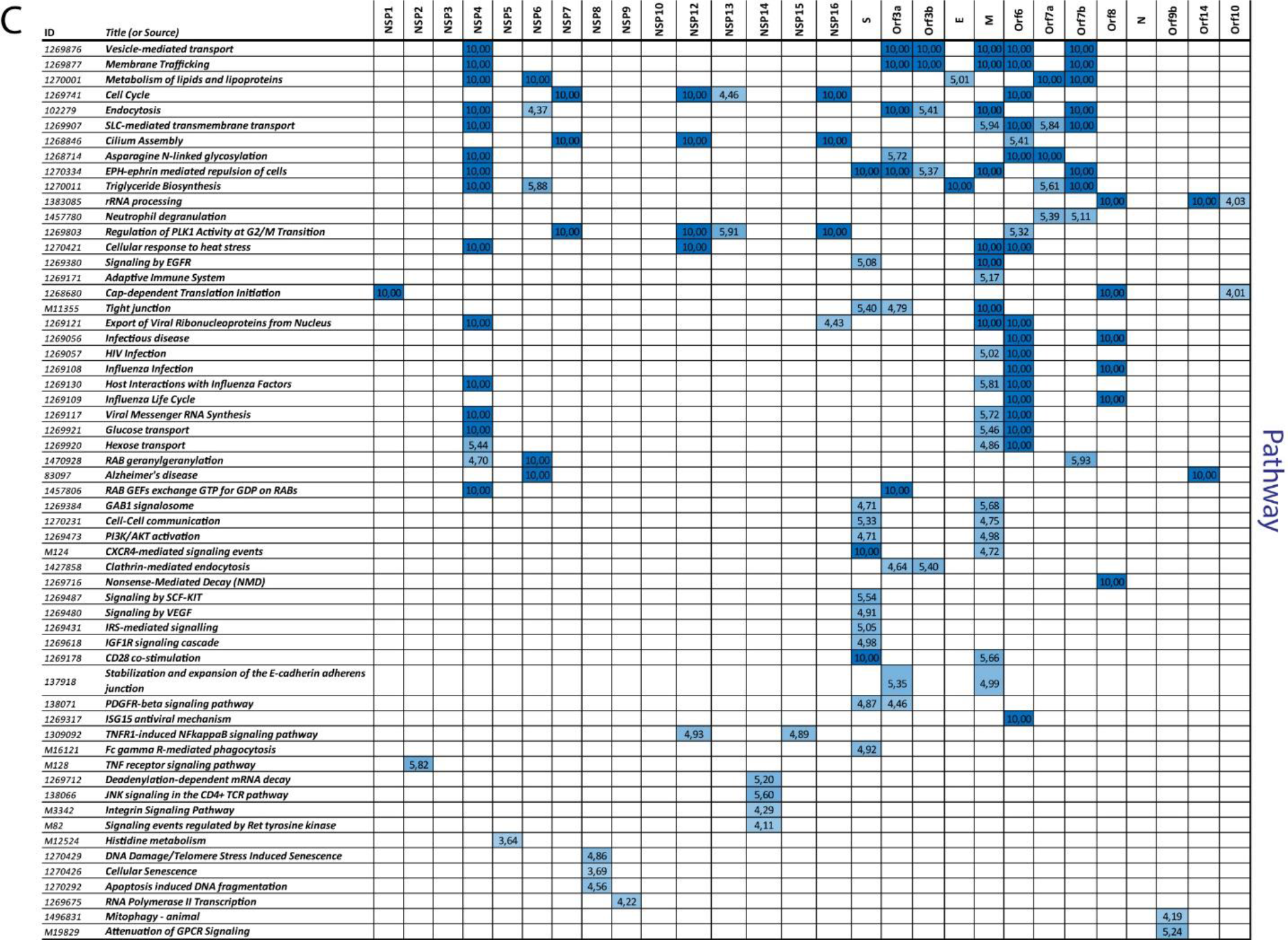
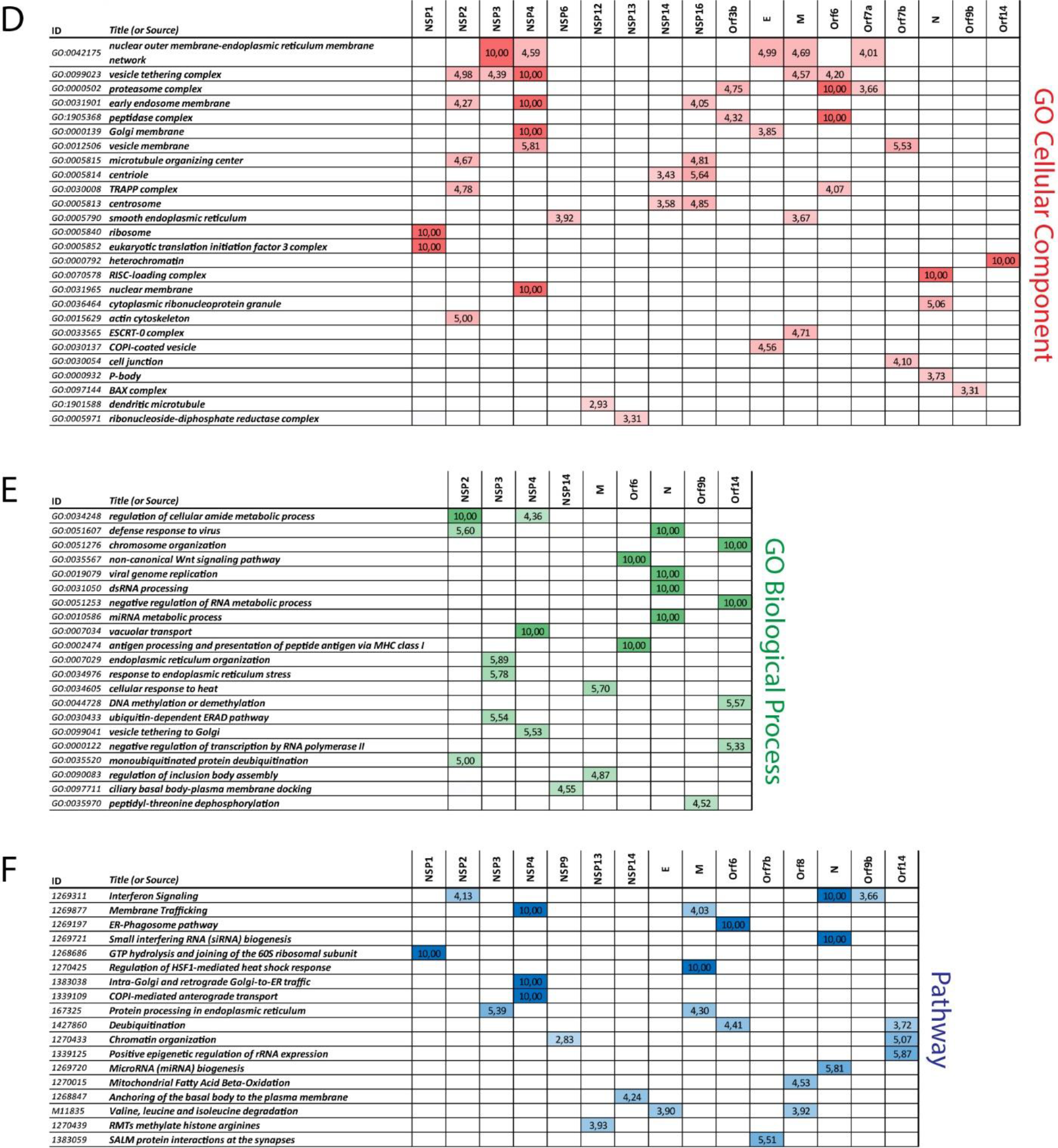
Comparative enrichment. Selected categories from ToppCluster analysis (see supplemental methods). **A**, **B** and **C** correspond to proximal interactome enriched categories detected in basal culture conditions. **D**, **E** and **F** are those detected enriched when analyzing increased or gained interactors after poly(I:C) treatment. For all panels, the -log_10_ p-value are highlighted with color gradients (red for GO Cellular Component; green for GO Biological Process; and blue for Pathway).

## Supplemental Material and Method

### Cloning

SARS-CoV-2 viral proteins coding sequences were cloned into pcDNA5 FRT/TO N-ter and C-ter BirA*Flag vectors by Gateway cloning (as in^1^) using our collection of Gateway-compatible entry clones as templates^2^ (see **Supplemental table 4** for sequences). All clones were sequence verified before cotransfection with pOG44 in Flp-In™ T-REx™ HEK293 cells (Thermo Fisher Scientific).

### Cell lines and sample generation

BioID samples were prepared as in^3^. Briefly, Flp-In™ T-REx™ HEK293 cells were grown in Dulbecco’s Modified Eagle’s Medium (DMEM, Gibco) supplemented with 10% fetal bovine serum (FBS, Sigma-Aldrich), GlutaMAX™ and Penicillin-Streptomycin (1x). Using the Flp-In system (Invitrogen), Flp-In™ T-REx™ HEK293 stably expressing BirA*Flag or FlagBirA* alone (for control samples), or N- and C-terminally tagged viral bait proteins were generated co-transfecting pOG44 with each pcDNA5 FRT/TO BirA*Flag-viral bait protein sequence plasmid. After selection (DMEM + 10% FBS + P/S + 200 μg/ml hygromycin B), three independent replicates of two 150 cm2 plates of sub-confluent (60%) cells were incubated for 24 hrs in complete media supplemented with 1 μg/ml tetracycline (Sigma), 50 μM biotin (Thermo Fisher Scientific). Cells were collected and pelleted (300 x *g*, 3 min), washed twice with PBS, and dried pellets were snap frozen. Each cell pellet was resuspended in 5 ml of lysis buffer (50 mM Tris-HCl pH 7.5, 150 mM NaCl, 1 mM EDTA, 1 mM EGTA, 1% Triton X-100, 0.1% SDS, 1:500 protease inhibitor cocktail (Sigma-Aldrich), 1:1,000 Turbonuclease (BPS Bioscience) and incubated on an end-over-end rotator at 4°C for 1 hour, briefly sonicated to disrupt any visible aggregates, then centrifuged at 45,000 x g for 30 min at 4°C. Supernatant was transferred to a fresh 15 mL conical tube. 25 μl of packed, pre-equilibrated Streptavidin Ultralink Resin (Pierce) were added and the mixture incubated for 3 hours at 4°C with endover-end rotation. Beads were pelleted by centrifugation at 300 x *g* for 2 min and transferred with 1 mL of lysis buffer to a fresh Eppendorf tube. Beads were washed once with 1 mL of lysis buffer and twice with 1 mL of 50 mM ammonium bicarbonate (pH=8.3), then transferred in ammonium bicarbonate to a fresh centrifuge tube and washed two more times with 1 ml of ammonium bicarbonate buffer. Tryptic digestion was performed by incubating the beads with 1 μg MS-grade TPCK trypsin (Promega, Madison, WI) dissolved in 200 μl of 50 mM ammonium bicarbonate (pH 8.3) overnight at 37°C. The following morning, 0.5 μg MS-grade TPCK trypsin was added to the beads and incubated 2 additional hours at 37°C. Following centrifugation at 2,000 x g for 2 min, the supernatant was collected and transferred to a fresh Eppendorf tube. Two additional washes were performed with 150 μL of 50 mM ammonium bicarbonate and pooled with the first eluate. The sample was lyophilized and resuspended in buffer A (2% ACN 0.1% formic acid). 1/3^rd^ of each sample was analyzed per mass spectrometer run.

### BioID data acquisition

MS samples were prepared from three biological replicates of each bait protein fused either with a N-terminal or a C-terminal BirA*Flag epitope tag in basal condition or following poly(I:C) (Sigma-Aldrich; P1530) transfection at 2μg/mL for 5hrs using PolyJet reagent (Signagen) prior to tetracycline and biotin induction, and analyzed on a Thermo Q-Exactive mass spectrometer. Samples were separated by online reversed-phase chromatography using a Thermo Scientific Easy-nLC1000 system equipped with a Proxeon trap column (75 μm ID x 2 cm, 3 μm, Thermo Scientific,) and a C18 packed-tip column (Acclaim PepMap, 75 μm ID x 50 cm, 2 μm, Thermo Scientific). The digested peptides were separated using an increasing amount of acetonitrile in 0.1% formic acid from 2 to 30% for 2 hours at a flow rate of 300 nL/min. A voltage of 2.4 kV was applied by the liquid junction to electrospray the eluent using the nanospray source. A high-resolution mass spectrometer Q-Exactive™ Thermo Scientific™ was coupled to the chromatography system to acquire the 10 most intense ions of MS1 analysis (Top 10) in data dependent mode. The MS analyses were performed in positive mode at a resolving power of 70,000 FWHM, using an automatic gain control target of 3e^6^, the default charge state was set at 2 and a maximum injection time at 120 ms. For full scan MS, the scan range was set between m/z 300 to 1600. For ddMS^2^, the scan range was between m/z 200 to 2000, 1 microscan was acquired at 17,500 FWHM, an AGC was set at 5e^4^ ions and an isolation window of m/z 4,0 was used.

### BioID data analysis

The proteins were identified by comparing all MS/MS data with the Homo sapiens proteome database (Uniprot, release March 2020, Canonical+Isoforms, comprising 42,360 entries + viral bait protein sequences added manually), using the MaxQuant software version 1.5.8.3. The digestion parameters were defined using trypsin with 2 maximum missed cleavages. The oxidation of methionine and N-terminal protein acetylation were defined as variable modifications. The Label-free quantification (LFQ) was done keeping the default parameters of the software. As for initial mass tolerance, 6 ppm was selected for MS mode, and 20 ppm was set for fragmentation data to match MS/MS tolerance. The identification parameters of the proteins and peptides were performed with a false discovery rate (FDR) at 1%, and a minimum of 2 unique peptides per protein. The LFQ values from the 30 control runs (regrouping FlagBirA* and BirA*Flag alone samples, from stable and transiently transfected cell lines, and w/wo poly(I:C) pretreatment) were collapsed to the three highest values for each given ID. These three values were defined as the control group for comparison with viral bait proteins triplicates. The statistical analysis was done by Perseus software (version 1.6.2.3). Briefly, the LFQ intensity of each sample were downloaded in Perseus and the data matrix was filtered by removing the potential contaminants, reverse and only identified by site. The data were then transformed using the log2(x) function. Before statistical analysis, 84 groups (28 bait proteins, N-ter, C-ter and poly(I:C) for each) were defined with 3 replicates per group. Only preys with detected values in all three replicates of a given viral bait protein were kept for further analysis. Missing values were then replaced from normal distribution separately for each column. Two-sample Student’s T-test was then performed comparing all three biological replicates of each bait and condition against the three controls runs. High confidence proximal interactors were defined by permutation-based FDR with a cut-off of 0.01. Perseus output with all experimental values is reported in **Supplemental Table 1**.

### BioID data analysis

The matrix (**Supplemental Table 1**) shows the average log_2_ fold change against control and the corresponding p- and q-values for each bait and condition. The poly(I:C) data were also compared against control (green font when significant) and against the corresponding condition (N-terminal tag for all viral bait proteins, except for NSP1). The poly(I:C) column indicates the status of each interaction when compared to the basal condition. ‘Gained’ and ‘Lost’ are self-explanatory, and ‘Decreased’ and ‘Increased’ were defined when the log2 fc against the basal condition was <(−1) or >(+1), respectively. ‘Unchanged’ corresponds to log2fc values between −1 and +1, and empty cells depict interactors not detected in the basal and poly(I:C) condition. The InDegree column depicts the number of bait proteins detecting a given interactor, regardless of the condition (N-ter, C-ter or poly(I:C)). This criterion was chosen to filter out the most connected interactors (8+), likely to be organelle-specific background not filtered using the BirA* alone control samples.

### Data annotation

Reported interactions/interactors were annotated using the ‘Reported Corona interactors’ table (**Supplemental Table 1**). In the ‘High conf. (7 or less baits)’ tab, the reported high confidence proximal interactions correspond to the (bait-prey) pairs identified either in SARS-CoV1/2 or other coronaviruses in previous studies (as indicated). The reported interactors only report the preys which have already been associated with the indicated coronaviruses. The tabs ‘Enrichment basal condition’ and ‘Enrichment poly(I:C)’ have been generated entering the lists of high confidence proximal interactors of each viral bait protein in the ToppCluster online tool^4^ (https://toppcluster.cchmc.org/). Selected enriched categories for comparison across bait protein proximal interactomes are represented in **Supplemental Figure 1**. All interactors individual annotations are shown in **Supplemental Table 2**, which was generated using the Metascape annotation tool^5^ (https://metascape.org/). For the **Figures 3-31**, the Metascape Express Analysis report was used to automatically select the functional categories enriched for each given bait protein. The detail output of the functional enrichment analysis, including the gene list assigned to each category for each bait protein, are provided in **Supplemental Table 3**. Finally, each viral bait protein proximal interactome was manually analyzed through literature and databases curation, and the noticeable features are reported in the main text.

### Confocal microscopy

Each N-ter tagged viral protein expression was inducted by 1μg/mL tetracycline and 50μM biotin for 24 hrs (in basal or poly(I:C) conditions) in stably transfected Flp-In™ T-REx™ HEK293 cells. Briefly, cells were grown on poly-lysine coated coverslips, rinsed with PBS before fixation with 4% paraformaldehyde for 15 min at RT. After fixation they were washed 3 times with PBS 0.1% Triton X-100. Cells were then blocked in PBS 0.01% Triton, 5% bovine serum albumin (BSA) for 30 min at 4°C. Transfected cells were then incubated overnight at 4°C with monoclonal anti-FLAG M2 mouse antibody (dilution 1:2,000, Sigma-Aldrich). After 3 washes for 10 min with PBS 0.01% Triton X-100, cells were incubated for 1 h at 37 °C with the Alexa Fluor 647-conjugated secondary antibody donkey anti mouse IgG (H+L) (1:2,000, Thermo Fisher Scientific) and streptavidin-Alexa Fluor 488 (1:20,000; Invitrogen). Cells were rinsed with PBS and nuclei were stained with 1 μg/ml 4’,6-diamidino-2-phenylindole DAPI (Thermo Fisher Scientific) in PBS for 5 min. Finally, cells were washed, desalted and mounted using Dako Fluorescent Mounting Medium (Agilent Dako, Santa Clara, CA, USA). Acquisition was performed on a Zeiss LSM700 confocal microscope connected to a Zeiss Axiovert 200 M equipped with an EC Plan-Neofluar 40×/1.30 numerical aperture and an oil immersion objective (Carl Zeiss AG, Oberkochen, Germany). Processing of the images was performed using Zeiss Zen 2 software and assembled using Adobe Illustrator.

### Interactome network setup and visualization

The starting point for constructing the interactome network is the BioID high confidence PPI data in the form of a list of pairs of bait-target interactors. There is a total of 28 baits and 2598 targets (modulo isoforms), with 10198 interactions. Based on this data we construct a graph with interactors and interactions represented by the vertices (nodes) and edges (links) respectively of a directed graph.. In addition to the observed BioID interactions, we include 3512 extra edges in the graph, representing the known interactions between targets. Thus, the final graph has a total of 2626 vertices and 13710 edges.

To visualize this graph, our objective is to determine an optimum placement of the nodes in threedimensional space that would minimize edge crossings and reveal the interactome’s structure to the best possible extent. We employ a custom, multi-stage, force-directed graph layout algorithm in 3D, based largely on the algorithm outlined by Hu^6^. We modulate the strength of the attractive forces in the physical model with a factor given by the combination of the experimental enrichment ratio results according to 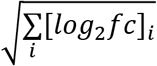, where i = N-ter, C-ter, poly(I:C), when present for the interaction. For interactions where this information is not available (such as the reported target-target interactions) we set this factor equal to 1. The node 3D coordinates obtained by minimizing the energy of the physical model are fed into code that creates a 3D visualization of the network by assigning a point to each node and a line connecting a pair of nodes when there is an interaction between the corresponding proteins. Our code is written in Python 3.8^7^ and makes use of several modules, primarily: NetworkX^8^ for graph operations, NumPy^9^ for array manipulation and numerical computations, pandas^10^ for data handling and Plotly^11^ for visualization.

### Online interactive web application

In order to make our results available to the scientific community in an usable and informative manner, we have implemented a web app accessible at http://www.sars-cov-2-interactome.org/, containing the interactive network visualization as well as multiple options for selecting and filtering the dataset to view subnetworks, providing the ability to focus on particular areas of interest and explore various levels of detail. In addition, we integrate and overlay information about the reported status of interactors and interactions with respect to SARS-CoV-1/2 or other coronaviruses, as well as the previously reported interactions between the targets (**Supplemental Table 1** for details, and main text references^1,2,3,17S^). The network customization options primarily consist of four layers: the first layer gives the ability to filter out targets when their total number of interactions with baits exceeds a configurable threshold, giving the ability to focus on targets of higher specificity in the interactome. At the same layer the target-target reported interactions can be turned on/off on the network. The second layer allows the user to select annotations of interest from three main dropdown categories (pathways, biological processes, cellular components). In the case of multiple selections, the annotations are combined, and the resulting network is the union of the individual annotated subnetworks. The third layer applies filtering to the selected network’s edges and nodes: these are based on the interaction’s poly(I:C) treatment result, the type of reported interaction, the interactors’ reported status, and a filter restricting to interactions of specific baits of choice. The various filtering criteria in this layer are combined by intersection, restricting the view to a portion of the nodes and edges of the previously selected subnetwork that satisfy them all. The last layer of customization refers to mostly cosmetic attributes of the network visualization, such as the ability to toggle text labels and edges, present a faded view of the full network in the background (thus giving a better global view of the location of nodes and edges in the interactome), and display 3D spherical shapes for the interactors (as opposed to flat 2D circular markers), better representing their physical size and actual location in space. We plan to continually improve the dashboard by adding useful features in future updates. Integrating with networks from other related studies in a comparative manner could provide an even more comprehensive resource that could yield important insights about the virus mechanisms of action. The dashboard web app was implemented as a Flask^12^ service, making use of the Dash^13^ module in Python 3.8.

## Supplemental Tables legends

**Supplemental Table 1. A.** High confidence proximal interactors detected by seven or less viral bait proteins. The number of baits detecting a given prey protein is indicated in the ‘Indegree’ column. The p- and q-values are extracted from the Perseus statistical analysis of MaxQuant data (Tab H for details). The ‘mean log2fc’ column represent the ratio between the detected preys in given bait assays against the control group. The basal condition ‘High confidence’ column indicates hits which are detected as high confidence proximal interactor in N-terminally and/or C-terminally tagged corresponding bait BioID. The ‘log2fc poly(I:C) vs. basal’ shows the log2 fold LFQ intensities changes between the preys detected after poly(I:C) treatment enrichment and the corresponding basal condition. Over 1 is considered as either enrichment or gain (if absent from the basal condition), between 1 and −1 corresponds to unchanged, and below −1 shows decreased or lost hits. When no value is indicated, it corresponds to interactions not detected in the poly(I:C) and the corresponding basal condition. The reported interactions columns display the bait-prey pairs previously identified in the literature (see tab **D** for details and references and main reference^178^). The reported interactors columns show preys which have been identified associated with coronaviruses in previous studies (regardless of the viral bait protein). **B.** Displays all interactions before indegree filtering. **C.** Shows the viral baits detection in the BoID experiments. Of note, most of the nondetected bait proteins are too short to generate enough or suitable tryptic peptide for MS detection. **D.** Reported interactions of all coronaviruses across the literature, with corresponding techniques and references. **E.** Comparative ToppCluster analysis (see Supplemental Method) output of all bait interactomes obtained in basal condition (union of N- and C-terminally tagged viral bait proteins BioID results). **F.** Input list for the analysis presented in E. **G.** Same as E, but with hits increased or gained after poly(I:C) treatment. **H.** Input enriched or gained interactors after poly(I:C) for the ToppCluster analysis presented in G. **I.** Raw output of the Perseus analysis of MaxQuant LFQ results. Columns G-JA: log2 transformed LFQ intensities for all biological replicates and conditions; Columns JB-MG: High confidence proximal interactors with an FDR<0.01 (’+’). Columns MI-MS: number of peptides detected for each prey protein and statistical significance. Columns MT to the end: statistical analysis of each bait-prey interaction.

**Supplemental Table 2.** Metascape custom annotation of all high confidence proximal interactors. The cell C1 is a text request which will display in the column C the interactors for which the entered term is present in any field of their descriptions.

**Supplemental Table 3.** Metascape functional analysis of each prey interactor.

**Supplemental Table 4.** SARS-CoV-2 cloning sequences used in this study.

